# Particle-based phasor-FLIM-FRET resolves protein-protein interactions inside single viral particles

**DOI:** 10.1101/2023.05.31.543036

**Authors:** Quinten Coucke, Nagma Parveen, Guillermo Solís Fernández, Chen Qian, Johan Hofkens, Zeger Debyser, Jelle Hendrix

**Affiliations:** Molecular Imaging and Photonics Division, Department of Chemistry, KU Leuven, Leuven, Belgium; Department of Chemistry, Indian Institute of Technology Kanpur, India; UFIEC, National Institute of Health Carlos III, Majadahonda, Madrid, Spain; Department of Chemistry, Center for Nano Science (CENS), Center for Integrated Protein Science Munich (CIPSM) and Nanosystems Initiative Munich (NIM), Ludwig Maximilians-Universität München, Munich, Germany; Laboratory for Molecular Virology and Gene Therapy, Department of Pharmaceutical and Pharmacological Sciences, KU Leuven, Leuven, Belgium; Dynamic Bioimaging Lab, Advanced Optical Microscopy Centre and Biomedical Research Institute, Hasselt University, Hasselt, Belgium

**Keywords:** FRET, Phasor, Phasor-FLIM, FLIM, particle-based, HIV-1

## Abstract

Fluorescence lifetime imaging microscopy (FLIM) is a popular modality to create additional contrast in fluorescence images. By carefully analyzing pixel-based nanosecond lifetime patterns, FLIM allows studying complex molecular populations. At the single molecule or single particle level, however, image series often suffer from low signal intensities per pixel, rendering it difficult to quantitatively disentangle different lifetime species, such as during FRET analysis in the presence of a significant donor-only fraction. To address this problem, we combined particle localization with phasor-based FLIM analysis. Using simulations, we first showed that an average of ∼300 photons, spread over the different pixels encompassing single fluorescing particles and without background, is enough to determine a correct phasor signature (standard deviation <5% for a 4 ns lifetime). For immobilized single- or double-labeled dsDNA molecules, we next validated that particle-based phasor-FLIM-FRET readily allows estimating fluorescence lifetimes and FRET from single molecules. Thirdly, we applied particle-based phasor-FLIM-FRET to investigate protein-protein interactions in sub diffraction HIV-1 viral particles. To do this, we first quantitatively compared the fluorescence brightness, lifetime and photostability of different popular fluorescent protein-based FRET probes when genetically fused to the HIV-1 integrase enzyme (IN) in viral particles, and conclude that eGFP, mTurquoise2 and mScarlet perform best. Finally, for viral particles co-expressing FRET-donor/acceptor labeled IN, we determined the absolute FRET efficiency of IN oligomers. Available in a convenient open-source graphical user interface, we believe that particle-based phasor-FLIM-FRET is a promising tool to provide detailed insights in samples suffering from low overall signal intensities.

**Why it matters:** Phasor-FLIM is an extraordinarily popular tool for fluorescence lifetime imaging analysis. However, it remains susceptible for low signal intensities, operational challenges and therefore required informed users and a clear analysis understanding. In this work we developed a convenient all-graphical workflow for quantitative phasor-FLIM in heterogenous and low-signal samples and applied it to quantifying absolute FRET efficiencies from protein-protein interactions inside single viral particles. Moreover, containing a well-illustrated theoretical introduction to time-domain phasor-FLIM, our paper helps novice users to correctly implement phasor-FLIM in standard microscopy practice.

## Introduction

Fluorescence lifetime imaging microscopy (FLIM) exploits the excited state fluorescence lifetime of fluorophores to generate image contrast (1–3). For FLIM data recorded using pulsed lasers and photon counting (i.e., time-domain FLIM), pixel-based analysis is typically performed via (multi-component) fluorescence decay fitting (4, 5) or more recently, the phasor approach to time-domain FLIM (from here on referred to as phasor-FLIM) (6). Wide applications of phasor-FLIM, such as biosensors based on FRET (7–10), the autofluorescence signature of a sample (11–16) or investigating the nanoscale diversity of crystalline materials (17) have been extensively described. Additionally, new techniques sprouted using phasor principles such as spectral phasor focused on unmixing (18), phasor S-FLIM focused on species photophysics (19), and several others (20–22).

The wide applicability of FLIM comes with some limitations, the most notable of which is heterogeneity in FLIM data. Lifetime heterogeneity in FLIM data between individual pixels, in general, is extracted and utilized by a per-pixel lifetime analysis. Heterogeneity within one pixel, on the other hand, presents a complication for quantitative FLIM regardless of whether decay-fitting or phasor-transformation based analysis is used (23, 24). In these approaches, the unmixing of contributing species is executed in different manners, each with benefits and drawbacks. While global lifetime fitting can provide an analysis for the whole FLIM image, it requires critical user input such as the number of exponentials and the fitting boundaries. The phasor approach attempts the unmixing of contributing factors in a graphical way but equally requires user input such as the instrumental referencing, the autofluorescence contribution and the location of the pure species in the phasor plot (6, 15). Nonetheless, it is generally accepted that the phasor approach allows for easy identification of data clusters and species assignment combined with convenient data representation using the image-phasor reciprocity (25). You can select pixels of the image and analyze this subset in phasor, or inversely, select phasor points that will represent a subset of pixels.

Despite the relative ease of use, the application of phasor-FLIM has primarily focused on bright dyes, abundant label-free autofluorescence (such as nicotinamide adenine dinucleotide, NADH) or overexpressed fluorescent protein systems (8, 15, 26, 27). Lower photon yields, sub-pixel species heterogeneity and other experimental complications (e.g., fluorophore maturation, photobleaching) are common experimental complications that render quantitative analysis challenging (28, 29). Fluorescent protein-based biosensors, for example, generally offer high signal intensities due to strong subcellular expression (30–34). However, FLIM-based Förster resonance energy transfer (FRET) protein-protein interaction studies at physiological cellular concentrations, are often carried out at much weaker overall fluorescence signals. Methodologies that facilitate quantitative analysis under conditions of low photon budget and/or signal/noise, such as, e.g., the recently developed particle-based phasor-FLIM approach, are therefore highly desirable (35).

As a specific low-signal example, we focus here on fluorescent protein-based FLIM-FRET of homo-interactions of human immunodeficiency virus 1(HIV-1) integrase (IN) enzymes inside single viral particles. The sub-resolution HIV-1 particles can incorporate a limited number of fluorescent proteins/fluorescently-labelled IN molecules. As such, they present a challenge for quantitative FRET analysis because of the overall low signal intensities. We previously employed acceptor photobleaching intensity-based FRET studies on the HIV-1 and murine leukemia virus (MLV) IN enzyme to show that the multimerization state of IN functionally changes during nuclear entry or after drug treatment of infected cells (36–38). However, since the IN-labelled donor (D) and acceptor (A) could oligomerize in different combinations in the FRET system used, intensity-derived FRET values are significantly underestimated by the presence of D-D homodimers, which in turn limited the overall FRET dynamic range. Furthermore, the intensity-based FRET analyses used was completely blind to possible sub-particle species heterogeneity.

In this paper we explored the grouping effect of pixels from a single particle. In particular, we look at what advantages it presents in the analysis of dim photon limited samples in which normal pixel-based phasor-FLIM analysis struggles. Additionally, we present the phasor-FLIM analysis in conjunction with the phasor theory to clarify phasor-FLIM for novice users. In doing so we present a fair evaluation of FRET data on the oligomerization of HIV-1 integrase.

## Theory

### Phasor approach to FLIM

The phasor plot representation of FLIM data was originally developed for frequency-domain FLIM analysis (6, 39–41). Frequency-domain FLIM utilizes a modulated excitation source at an angular frequency 𝜔. The frequency-dependent demodulation 𝑀_𝜔_ and phase shift 𝜑_𝜔_ are characteristics of the emitted fluorescence, and each quantity can be related to the fluorescence lifetime:

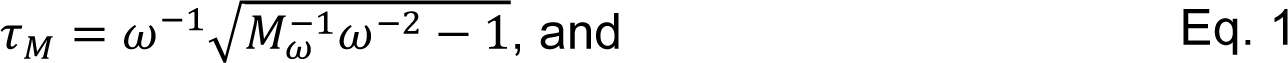

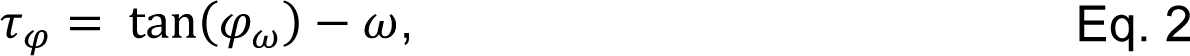

where the phase (𝜏_𝜑_) and modulation (𝜏_𝑀_) lifetimes are equal for high-signal and pure-species data (see also Supplemental Figure S8). Next to this, every lifetime can also be represented in a polar plot as a vector with length equal to the demodulation 𝑀_𝜔_and an angle with respect to the abscissa equal to 𝜑_𝜔_. This vector is called the phasor, a portmanteau of phase vector. The point determined by the phasor is described using the Cartesian coordinates 𝑔_𝜔_ and, 𝑠_𝜔_ which are the first cosine and sine Fourier coefficients, respectively, of the time-dependent fluorescence signal 𝐼(𝑡) (Supplemental Figure S1A):

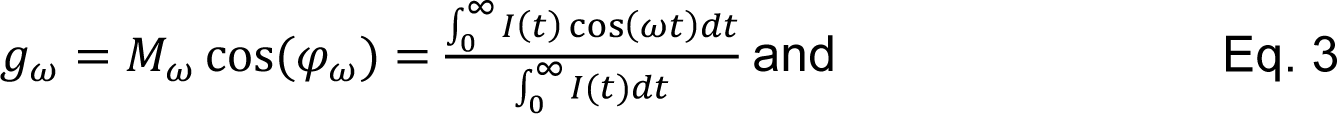

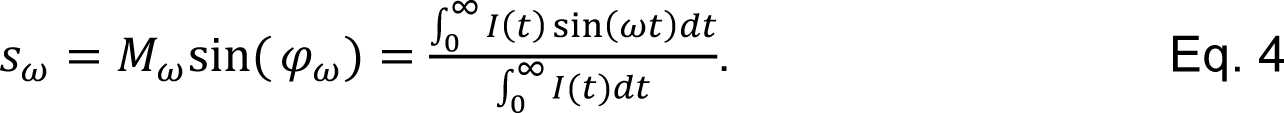

A direct relation between 𝜑_𝜔_ and 𝑀_𝜔_ is found when investigating a mono-exponential decay, where Eqs. 1 and 2 result in the same lifetime and can be equaled:

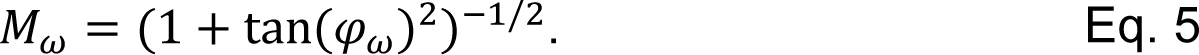

𝑀_𝜔_ and 𝜑_𝜔_ can subsequently be written as:

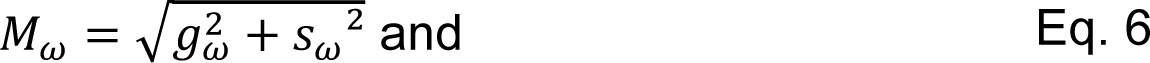

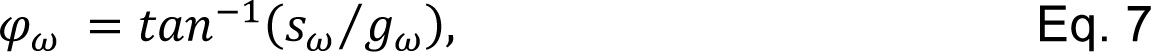

which allows rewriting Eq. 5 in the Cartesian coordinates of the polar plot:

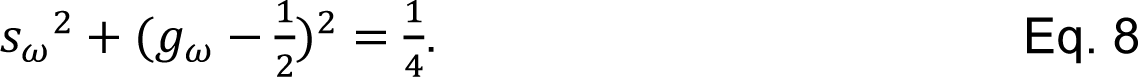

Eq. 8 describes the so-called universal semi-circle, on which all mono-exponential signals will be spread out, with the phasors for shorter lifetimes lying close to the point (1,0), and larger lifetimes closer to (0,0) at the origin point of the semicircle (see also Supplemental Figure S9). Combining Eq. 8 with Eqs. 3 and 4 subsequently allows calculating the 𝑔 and 𝑠 coordinates using the angular frequency and the monoexponential lifetime.

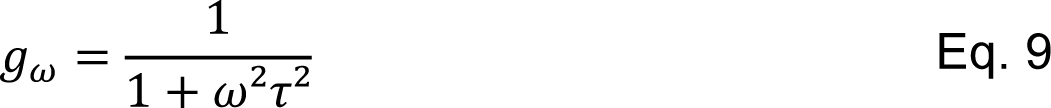

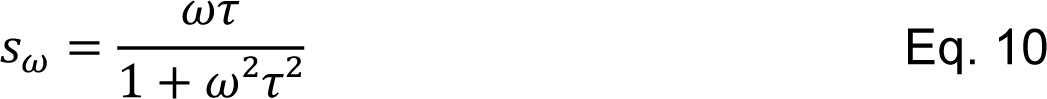

All multi-exponentials, which are a sum of mono-exponentials, are linear vector combinations of the constituent phasors, and thus will be found inside the described semi-circle. For a combination of 𝑃 species the resulting phasor coordinate is calculated by the sum of fractional photon contributions (𝑓_𝑖_) of every pure species *g*_𝜔,𝑖_, *S*_𝜔,𝑖_ (see Supplemental Figure S1A and S1E):

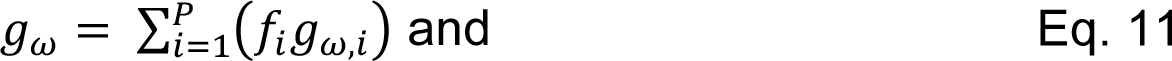

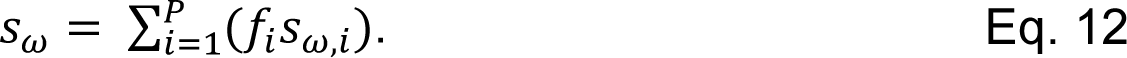

### Phasor FLIM using pulsed excitation

When using pulsed excitation and time-correlated single photon counting (TCSPC) detection, as in time-domain FLIM, 𝜔 is no longer the modulation frequency but is rather determined by the TCSPC range (𝑇, in units of time), i.e., the total time period during which detected photons are timed. In this case, the angular frequency 𝜔_𝑇_ used for phasor transformations is given by:

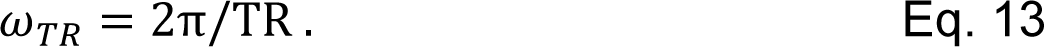

When calculating the phasor for a pixel, or by extension, an image, the idealized scenario where mono-exponential components are directly positioned on the semicircle does not hold true. Most importantly, the fluorescence decay and TCSPC range do not start at the same point relative to the TR, and even for a pure species, its mathematically mono-exponential decay will be convoluted with the instrument response function (IRF) of the measuring device, both effects significantly throwing off any phasor determination. Therefore, a reference measurement is necessary of a stable, well characterized and mono-exponentially decaying dye. Alternatively, the pure IRF signal can be used, representing a lifetime of 0. The reference’s known fluorescence lifetime 𝜏_𝑟_ is first used to calculate (using Eqs. 6,7,9,10) the expected demodulation 𝑀_𝑟_ and phase shift 𝜑_𝑟_ values for the reference (see Supplemental Figure S1B)

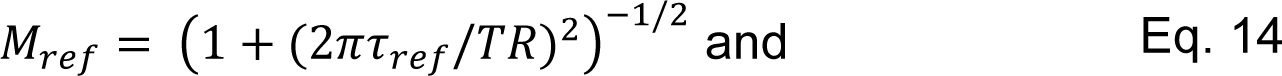

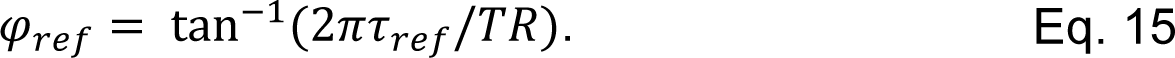

These values are subsequently used to calculate the phasor values for the instrument:

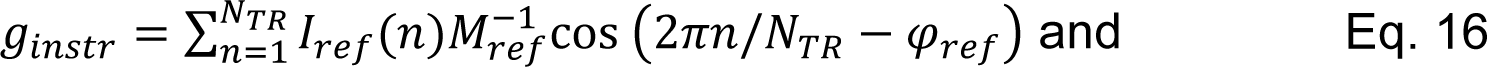

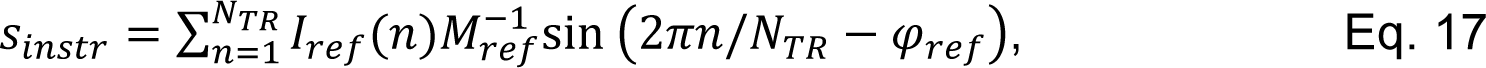

where 𝑁_𝑇_ is the total number of time channels in the TCSPC range and 𝐼_𝑟_(𝑡) the normalized intensity of the reference in timebin 𝑡. These instrumental phasor coordinates can subsequently be used to calculate the phase and demodulation of the instrument, 𝜑_𝑖𝑡_ and 𝑀_𝑖𝑡_ via Eqs. 6 and 7. This finally allows to display instrument-dependent time-domain data in an instrument-independent polar plot:

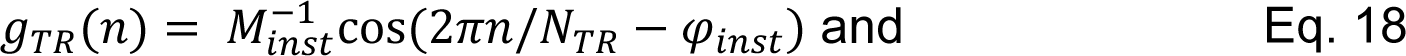

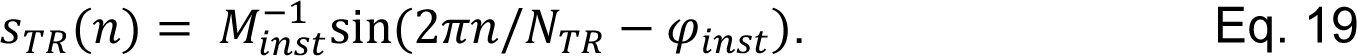

Every photon is assigned such coordinates depending on its arrival time channel in the TCSPC range, and depending on the subsequent analysis, photon phasors were either averaged per pixel or per identified object. An overview figure of time-domain FLIM-phasor illustrating pulse frequency impact and reference implementation is provided as Supplemental Figures S1B-D and Supplemental figure S9.

### Resolving mixtures of pure species

Pixels containing photons that originate from a single type of pure (mono-exponentially decaying) fluorophore translate to phasor values that lie on the semicircle. Phasors from pixels containing a mixture of two pure fluorescent species, with each having a different fluorescence lifetime, on the other hand, are located on a straight line connecting the two constituent pure phasors (Eqs. 11 and 12). The position of the mixture‘s phasor is determined by both their fluorescence lifetime and relative intensity (brightness) of each species, which is in turn defined by their relative concentration. It is possible to disentangle both. In the case of a mixture of an unquenched and FRET-quenched fluorophore, for example, the concentration fraction 𝑓_𝐶_ of each species is determined from the lifetimes and the intensity fraction 𝑓 as follows:

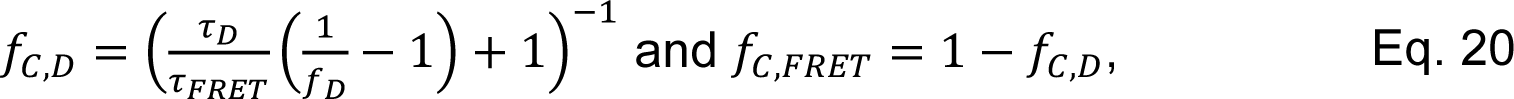

where 𝜏_𝐷_ and 𝜏_*FRET*_ are the Donor-only and FRET-quenched lifetimes, resp., and 𝑓_𝐷_ is the intensity fraction of the D-only species. For the derivation of this formula, the reader is referred to Supplemental Note SN1. Practically, the fraction line is manually drawn, starting from the phasor value of the D-only species and through the center of mass of the mixture’s phasor value. This then renders the fractions and the lifetime of the FRET species.

### Phasors in the presence of background signal

Background signals (autofluorescence, detector dark counts, scatter, water Raman, laser reflections…) will cause the phasor value of a pure species to be shifted from the semicircle, preventing correct lifetimes to be estimated. By measuring the phasor signature of the background (BG) in a reference experiment, however, it can be used to draw a fraction line through the phasor of the unknown pure species, whose lifetime can then be correctly read at the intersection of the fraction line and semicircle. For FRET experiments, the BG fraction is estimated by the position of measured donor-only sample.

### Quenching trajectory of pure FRET species

In the case of a pure fluorophore that is quenched by FRET, its phasor describes a trajectory starting from the D-only species (no FRET) along the semicircle to approximate a 0-ns lifetime (100% FRET) as described by the resulting mono exponential lifetimes on the semi-circle by the formula:

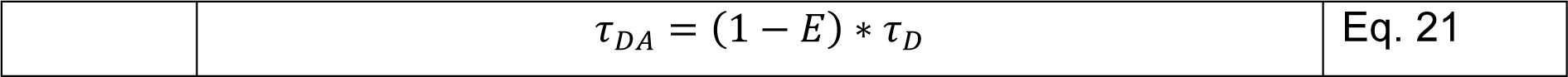

(Fig. S1F). In reality, however, background (𝐵) and a possible contribution from donor molecules with no nearby acceptor (𝐷) will cause this trajectory to deviate from the semicircle. For points resembling higher FRET states the fluorescence quantum yield of the involved species decreases leading to increased fluorescence contribution of 𝐷 and 𝐵. Taking this into consideration, the quenching trajectory in a non-perfect condition folds back onto itself and ends at the phasor determined by the remaining 𝐷 and 𝐵 (Fig. S1G). The quenching trajectory connecting all possible FRET efficiency 𝐸𝐸 states, can therefore be determined by fraction lines between all monoexponential points on the semicircle < 𝜏_𝐷_ and the *BG* & *D_only_* phasor. *g* and s coordinates of the quenching line are given by:

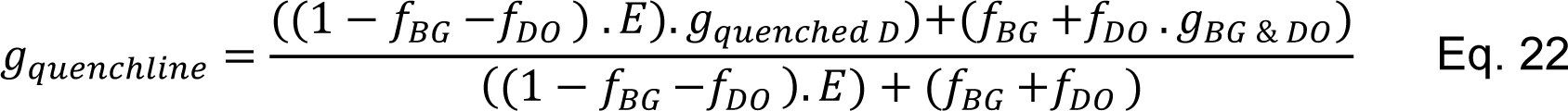

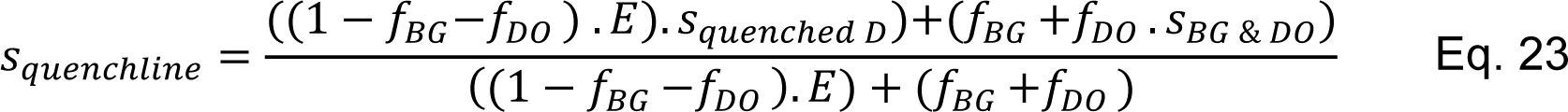

in which 𝑔_*quenched D*_ and 𝑠_*quenched D*_, the coordinates of the pure quenched form on the semicircle, can be found using Eqs 9 and 10 in which the lifetime of the quenched form is 𝜏_𝐷 𝐷𝑖𝑜_(1 − 𝐸𝐸). From the formulation it is clear that with increasing FRET of the pure quenched state the fractional contribution of the 𝐵 & 𝐷 phasor becomes larger. Construction of the quenching line and its contributing factors are displayed in supplemental Figure S1 F-I.

## Materials and Methods

### Analysis software

All analysis of imaging data (simulated or experimental .ptu files) was done in the open source Pulsed Interleaved Excitation (PIE) Analysis with MATLAB (PAM) software, a versatile software package offering a variety of analysis tools, including phasor analysis of FLIM data (42). The software is available as a source code, requiring MATLAB to run, or as compiled standalone software compatible with Windows or MacOS at http://www.cup.uni-muenchen.de/pc/lamb/software/pam.html or hosted in Git repositories under http://www.gitlab.com/PAM-PIE/PAM and http://www.gitlab.com/PAM-PIE/PAMcompiled. Sample data is provided under http://www.gitlab.com/PAM-PIE/PAM-sampledata. A detailed manual is found under http://pam.readthedocs.io. The workflow of the software as applied in this paper is described in Supplemental Figure S2. Simulated photon data was generated by a subsection of PAM. This module is illustrated in Supplemental Figure S3. Simulation input parameters included: TCSPC range (50 ns), pixel intensity (defined by the number of frames pixel dwell time and the particle brightness *ε*), fluorescence lifetime, background intensity (0 kHz), diffusion coefficient *D*, number of species and their concentration, simulation box size, pixel size (50 nm) and image size, and (Gaussian) IRF width (250 ps). For the single-particle simulations, a single species of *D* = 0 µm²/s was used. For the concentration fraction line simulations, *D* = 100 µm²/s for both species and the fluorescence lifetime and brightness of one species was four times larger than that of the other species (resp. 4 ns and 1 ns), mimicking a species and its 75% quenched form. Particle detection was performed using the *Particle Detection* functionality within PAM, which localizes particles in the fluorescence intensity images using eccentricity (0.5), counts (>300), size in pixels (min:15, max 100) and a wavelet depth of 3, using the simple wavelet method (35). Colocalization of dsDNA particles was performed using the centroid positions of particles in the FRET donor and directly excited FRET acceptor images as input via a nearest neighbor search with a maximum distance tolerance of 5 pixels for successful colocalization. Statistical analysis was performed using © 2022 GraphPad Software. For particle intensity tests and comparison of photobleaching, a Shapiro-Wilk tests was first used to check normality of data. When no normality was found, a non-parametric (one-way ANOVA, Kruskal-Wallis) test was used to compare datasets of each IN-FP.

### Determination of a phasor cloud center position

The shape of a given phasor ‘cloud’ at low signal intensities is determined by inherent shot noise on the one hand, and the spectroscopic contribution of the different emissive species, i.e., the (autofluorescence) background and the different lifetime species in the observed spectral window. To determine the center position, we calculated a ‘center of mass’, where the average phasor location of all pixels within the phasor cloud is determined, with each pixel weighed for the photon content of that point.

### Background and donor only fraction determination

Background samples are measured specifically per experimental case and aim to determine the contributing component of background, stray light, dark counts and autofluorescence. The phasor analysis is performed identical to a normal sample analysis to determine the resulting background phasor location. From all resulting phasor positions the photon weighted average position is determined and used as general background phasor. For DNA measurements the prepared non-labeled surface functionalized with PEG is imaged. In case of virus measurements, the viral particle itself is the main source of background and autofluorescence. Therefore, a photobleached region is imaged where viral particles are present in the scanned frame. The donor only contribution in a FRET experiment skew the phasor to the pure donor phasor position. Most often this value is not determined.

### DNA strand hybridization

All DNA strands were purchased from IBA LifeSciences (Göttingen, Germany). For the single-color dsDNA experiments the sense strand that was used is 5’-GGCTC GCCTG TGTXG TGTTG TATGA TGTAT TCGGC AGTGC GGG-Biotin in which the X marks the 14^th^ position that is labeled with ATTO 488 and biotin-labeled at its 3’-end. A compatible unlabeled antisense strand was used (5’-CCCGC ACTGC CGAAT ACATC ATACA ACACA ACACA GGCGA GCC) for annealing into a single labeled dsDNA. For our double labeled dsDNA, the antisense strand (5’-Biotin-TTTTT AAGTT TGTGA TAGTT TGGAC TGGTT YGTGA AGAAA AZCGC CGAAA A, with Y and Z an Alexa Fluor 488 and ATTO 647N label respectively) is used covalently labeled with Alexa Fluor 488 on position 31 followed by an ATTO647N label 11 nucleotides further. In addition, the antisense strands are labeled with biotin on the 5’ end. The complementary sense strand (5’-TTTTC GGCGA TTTTC TTCAC AAACC AGTCC AAACT ATCAC AAACT TAAAA A) is unlabeled. Sense and antisense DNA strands were hybridized by centrifuging the lyophilized strands for 1 min at 1000 g. Next, phosphate buffered saline (PBS, Merck, 806552, Sigma-Aldrich, Darmstadt, Germany) was used to dilute to a final concentration of 100 µM. The solution was homogenized by resuspension and vortexing. Afterwards, the top and bottom strand were annealed using a PCR machine (Doppio, VWR®-thermocyclerseries, VWR, part of Avantor, Radnor, US) increasing the temperature to 95 °C (3 min) and cooling down from 85 to 4 °C at a 1 min per degree to allow specific hybridization towards a final concentration of 10μM. Finally, the PCR product was transferred to a -80°C Eppendorf (Catalog Nr. 0030125215, Eppendorf Belgium) and directly flash-frozen in liquid nitrogen. Afterwards, the product was stored at -80°C. In order to remove the excess free dye as revealed by fluorescence correlation spectroscopy, a PD-10 desalting column was used.

### DNA experiments

The glass surface immobilization protocol involved surface cleaning and surface functionalization. Firstly, the chamber glass coverslips (#1.0 chambered coverglass, Lab-Tek Cat. No. 155411) was cleaned by an incubation with 10 mM SDS for 1 hour at room temperature. Next, the surface was washed several times with *Milli-Q* followed by surface activation with UV-Ozone for 15 minutes. Functionalization was performed by adding 10 μg/mL of the passivation agent (PLL-PEG, SuSoS, Dübendorf Switzerland) together with PLL-biotin-PEG in a 1:1 ratio and left to incubate at room temperature for 30 minutes. After this, the surface was rinsed three times with DNA resuspension buffer (150 mM STE-buffer, BP2478-1, ThermoFisher scientific, Geel, Belgium) without letting the surface dry out. This was applied after each incubation step of the functionalization. Once the surface contained biotin, 10μg/mL Neutravidin (ThermoFisher scientific) was added and incubated for 30 minutes followed by another three wash steps with 150 mM STE-buffer. Finally, DNA was diluted to the picomolar range using 150 mM STE-buffer and incubated for 30 minutes after which excess DNA was washed away.

### Microscope

The microscope presented in this work is a custom-built confocal microscope. Its schematic is shown in Supplemental Figure S4. The base microscope is an Olympus IX 70 modified with external pulsed excitation lasers, a scan unit and multichannel single photon detection. Five laser lines are available of which 4 are diode pulsed lasers (LDH series, PicoQuant GmbH, Berlin, Germany) and one supercontinuum laser (Solea, PicoQuant GmbH, Berlin, Germany). Laser pulsing and synchronization was set in the acquisitioning software SymphoTime 64 ‘1+2’ (PicoQuant GmbH, Berlin, Germany). A multichannel diode laser driver (PDL 828 Sepia2, Picoquant) controls the laser frequency at 20 MHz to ensure full fluorescence decay when measuring donor and acceptor fluorophores with pulsed interleaved excitation. (Dichroic) Mirrors guide the laser lines into a polarization-maintaining single mode optical fiber (PMC-400Si-2.6-NA012-3-APB-150-P, Schäfter+Kirchhoff GmbH, Hamburg, Germany) via a lens-based coupler (60FC-4-RGBV11-47, SuK). Light is collimated again via a lens-based collimator (60FC-L-4-RGBV11-47, SuK). Excitation light is directed at a dichroic quadband mirror (zt405/488/561/640rpc or zt440/510/561/640rpc depending on needed excitation line, AHF, Tübingen-Pfrondorf, Germany) held in place by a kinematic fluorescence filter cube (DFM1/M, 30 mm cage compatible, Thorlabs, Munich, Germany) for easy optics switching, and reflecting the excitation beam to the scanhead (TILL Yanus IV digital scanner, FEI Munich, Gräfelfing, Germany) mounted straight onto the backport of the IX70. Scan motion is controlled with in-house software in C# Microsoft Visual Studio in combination with a National Instruments box (USB-6361 Multifunction I/O Device, NI, Austin, USA) that steers the xy galvo axis via a TILL photonics Scan Control Unit (SCU, FEI Munich, Gräfelfing, Germany). Pixel dimensions, number and dwell time are controlled by this software. Upon transmission of emission light through the quadband dichroic, the light is focused (150 mm AC254-15-A-ML, Thorlabs) onto a 50-µm pinhole (PS50S, Thorlabs), after being collimated again by (50 mm AC254-050-A-ML, Thorlabs). Detection is arranged over 3 detection channels, splitting the bundle with dichroic mirrors. For DNA experiments, both the 485-nm and 640-nm laser diodes (LDH-D-C-485 and LDH-D-C-640, Picoquant, Berlin, Germany) were used in pulsed interleaved excitation mode using the zt405/488/561/640rpc quadband. Emission light is split on a 560-nm-longpass (H560LPXR, AHF). Reflected light is cleaned up by a 530/50m (HQ, AHF) and recorded on an avalanche photo diode (APD, τ-SPAD, PicoQuant). Passing light is cleaned up by a 705/100 (ET bandpass, AHF analysentechnik, Tübingen-Pfrondorf, Germany). For mNeongreen, mClover3 and eGFP the 485-nm excitation source was used (LDH-D-C-485, Picoquant, Berlin, Germany) and emission detected identically to the ATTO 488 labeled DNA. mTFP1 and mTurquoise2 were excited using a 440-nm diode (LDH-D-C-440, Picoquant, Berlin, Germany) in combination with the zt440/510/561/640rpc quandband BS. Emission light was reflected on the 560 nm longpass and detected by an APD (τ-SPAD, PicoQuant) after passing a cleanup filter (480/40, Brightline HC, AHF analysentechnik, Tübingen-Pfrondorf, Germany). mVenus was excited using a 510nm diode (LDH-D-C-510, Picoquant, Berlin, Germany) in combination with the zt440/510/561/640rpc quandband BS. Emission light was reflected on the 560-nm longpass and detected by an APD (τ-SPAD, PicoQuant) after passing a cleanup filter (540/15, Brightline HC, AHF analysentechnik, Tübingen-Pfrondorf, Germany). mCherry and mScarlet2 were excited using a 560-nm supercontinuum laser (Solea, Picoquant, Berlin, Germany) in combination with the zt440/510/561/640rpc quandband BS. Emission light was passed through the 560-nm longpass and detected by an APD (τ-SPAD, PicoQuant) after passing a cleanup filter (600/37, Brightline HC, AHF analysentechnik, Tübingen-Pfrondorf, Germany). For FRET experiments with mTurquoise2-mVenus all mission light of donor and acceptor is reflected on the 560-nm longpass and split on a 507-nm-longpass dichroic (H507LPXR, AHF analysentechnik, Tübingen-Pfrondorf, Germany) after which a 480/40 and 540/15 cleanup filter is used for mTurquoise2 and mVenus respectively before detection. In case of eGFP-mScarlet the eGFP emission is reflected on the 560-nm longpass is detected, after cleanup with a 530/50m filter. For mScarlet the emission passes the 560-nm longpass and is cleaned up with a 600/37 filter.

APD detectors were powered by a dedicated power supply (DSN-102, PicoQuant). APD NIM signals are directed towards the HydraHarp 400 (Picoquant) to supply it with photon timing information. For all data acquired on the microscope a 60x water objective (Olympus UPlanSApo 60x/1.20 W Ꝏ/0.13-0.21/FN26.5) was used. The microscope was positioned on a vibration free isolated optical table (S-2000 series Stabilizer^TM^, Newport Spectra-Physics BV, Utrecht, Netherlands).

Frequent system checks, setup alignment and quantifying confocal volume parameters were performed with fluorescence correlation spectroscopy (FCS) using ATTO425-COOH, ATTO488-COOH, Alexa Fluor 546 or ATTO655-COOH (ATTO-TEC GmbH, Siegen, Germany) (Alexa Fluor™, Thermofisher Scientific). Data was analyzed using PAM-*FCSfit* (Supplemental Figure S5A). As references for phasor-FLIM, organic dyes were measured using the same optical setup for the assigned detection channel. Reference lifetimes were experimentally determined via PAM-*Taufit* (Supplemental Figure S5B) where the decay was fit using a reconvolution fit for a mono-exponential with loaded IRF. Dyes used were ATTO488-COOH, ATTO425-COOH, Alexa Fluor 546 and ATTO 647N-maleimide.

### Fluorescent HIV-1 particles

HIV-1-derived viral particles (Vesicular stomatitis virus glycoprotein (VSV-G)-pseudotyped) with fluorescent protein labeled integrase were synthesized using Vpr-mediated trans-incorporation (43, 44). HEK293T cells (6.5×10^6^) were seeded in 10 cm petri dishes with DMEM supplemented at 2% FBS and 50 μg/mL gentamicin (Invitrogen). For transfection medium was replaced with Opti-MEM® (Life Technologies, ThermoFisher Scientific) supplemented with gentamicin (Invitrogen). Cells were transfected with branched polyethylenimine (bPEI, 10 μM stock solution, Merck, Sigma-Aldrich, Darmstadt, Germany) when 90% confluency was reached. A three-plasmid system, consisting of 5 μg pVSV-G, 15 μg pNL4–3. Luc.R−.E− and Vpr-IN-FP-encoding plasmid (5 µg for single FP viruses, twice 2.5 µg for viruses containing two fluorescent proteins) was used in transfection. pVpr-IN-FP constructs with mTFP1, mTurquoise2, mNeongreen, mClover3, mScarlet, mRuby3, eGFP and mCherry were used. pVpr-IN-FP constructs for mTFP1 and mVenus are made according to Borrenberghs et al. (44). Other pVpr-IN-FP constructs were made by cloning the original pVpr-IN-eGFP plasmid introducing restriction sites via primers (43). Transfection lasted for 6 hours at 37°C and was terminated by replacing the medium with fresh 37°C Opti-MEM® (Life Technologies, ThermoFisher Scientific) supplemented with 50 μg/mL Gentamycin (Invitrogen). Virus supernatant was collected 48 h after initiating transfection, filtered through a 0.45-μm filter (Minisart® Syringe Filter, Sartorius, Göttingen, Germany), and concentrated by several washing steps in FBS-free Opti-MEM® and virus particles were concentrated by ultracentrifugation on a 60% (w/V) iodixanol cushion at 21°C (131,500 x g, 90 min, SW28 rotor, Beckman Coulter, Ireland) to a final volume of 1 mL. Iodixanol was removed by ultrafiltration (Vivaspin, MWCO 50K, Merck, Overijse, Belgium). All generated viral particles were kept up to 2 months at -80°C, without thaw-freeze cycles.

### Viral particle plating and glass coating

For imaging purposes, concentrated virus particles were freshly thawed and diluted in phosphate-buffered saline (PBS, Merck, 806552, Sigma-Aldrich, Darmstadt, Germany) based on the p24 antigen concentration in the supernatant, as determined by the p24-specific enzyme-linked immunosorbent assay (ELISA). Then, 200 μL of the virus dilution containing 1µg-3µg p24 antigen was transferred to a poly-D-lysine coated glass coverslip (#1 chambered coverglass, Nunc™ Lab-Tek™, Cat. No. 155411, Thermo Scientific) and incubated at 37 °C for 3-5 hours. The fixed virus particles were gently washed twice with PBS, after which 200 μL PBS was added to each well and the wells were sealed with parafilm to avoid evaporation. Poly-D-lysine coating was done by incubating the wells for 20 minutes at 37°C in a 0.4 mg/ml solution (4X) of poly-D-lysine (Merck, P7280-5MG, Sigma-Aldrich, Darmstadt, Germany), made from a 1 mg/ml stock, and followed by washing three times with PBS. Finally, all PBS was removed, and coverslips were air dried in a sterile flow for 30 min.

## Results

### Pixel binning improves phasor-FLIM of dim particles

As quality of phasor-FLIM depends on photon content in the fluorescence decay, we first quantified the effect of pixel binning on phasor-FLIM data. Specifically, we examined the effect of grouping pixels from the same sub-resolution object on the accuracy of the estimated lifetimes. Practically, we simulated confocal FLIM data of non-diffusing fluorophores exhibiting a 4-ns fluorescence lifetime. Because of their random axial location in the simulation box, the resulting image displayed particles with varying intensity in the image plane (Fig. 1 *A*).

**Figure 1.**
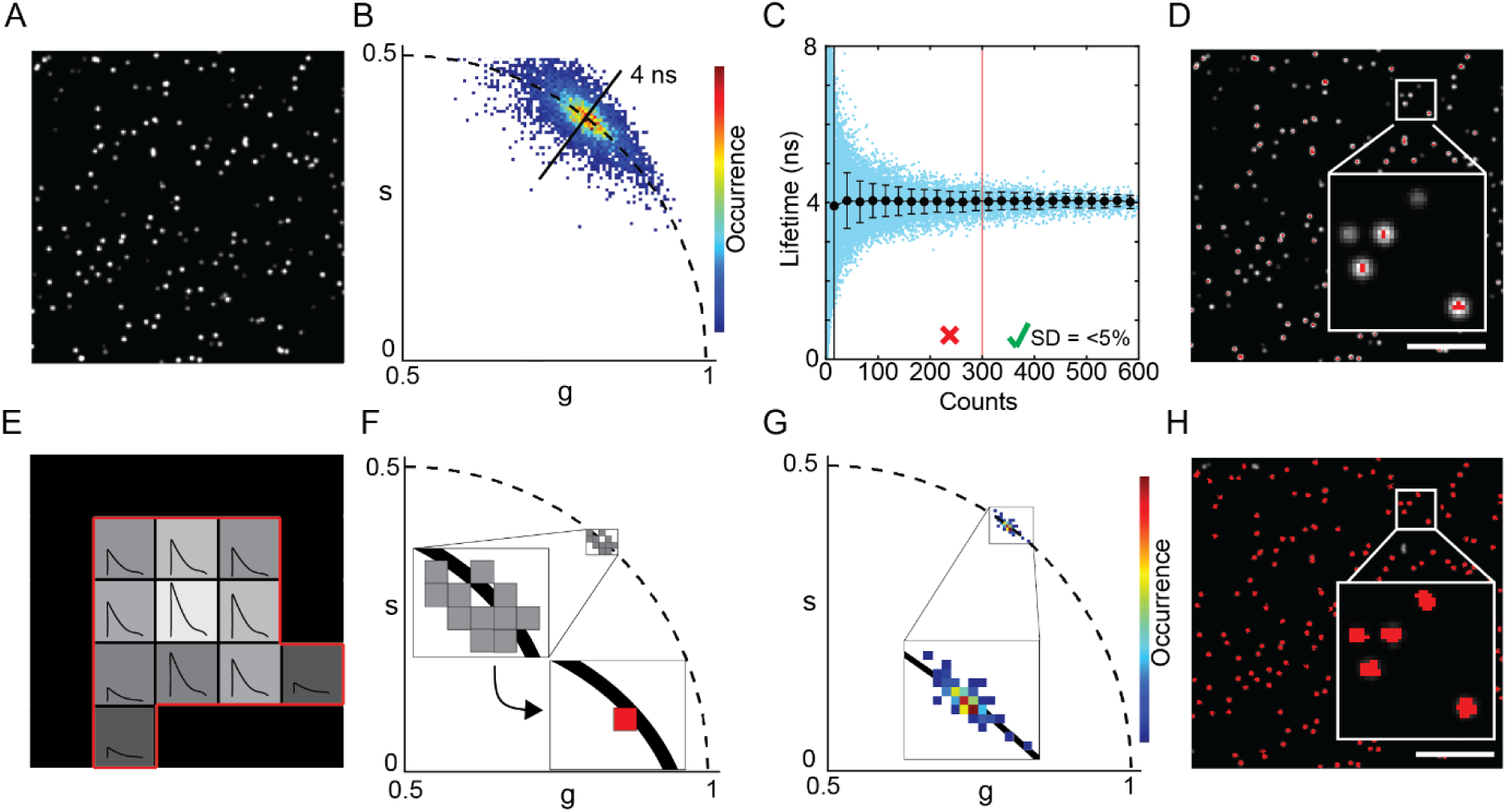
Pixel binning for particle phasor-FLIM: (A) Intensity image of simulated sub-resolution single particles with variety of intensities but a constant lifetime of 4 ns. (B) Phasor analysis of the simulated photon data results in a widespread on the phasor plot where every pixel (threshold for pixels with more than 5 counts) is assigned one phasor. (C) In blue: scatterplot of the pixel counts against the phase lifetime determined from the phasor plot. In black: standard deviation and mean lifetime per bin. Bins are defined as 25 counts wide, mean and standard deviation are plotted in the center coordinate of each bin. (D) Intensity image color coded in red if the pixel counts >300, inset is a 5×5 µm box. (E) Schematic of a single particle consisting of pixel from the same sub-resolution origin and showing identical decay lifetime composition but different amplitudes. (F) Pixel binning improves the particle phasor. (G) Simulated data from (A-D) analyzed in phasor with particle-based pixel binning shows a compact phasor distribution. (H) Particles color coded if the particle counts are >300 counts. All scalebars are 5 µm.

Due to the overall low intensity per pixel, the resulting phasor plot exhibited a relatively large data spread (Fig. 1 *B*). When plotting the phasor determined lifetime (average of phase and modulation lifetime, Eq. 1-2) for every pixel, we observed an increase in lifetime precision for increased photon counts of the pixel (Fig. 1 *C*). We determined that a count of 300 photons was ideally needed to obtain lifetime values within a 5% deviation range of the true value. This value can therefore be considered a lower limit, acknowledging that a higher photon count is advised for more complex phasor component deconvolutions. Using this 300-photon count threshold, we false-color-coded the intensity image for the pixels that meeting this set threshold and observed that relatively few pixels from the imaged particles were, in fact, colored (Fig. 1 *D*). Of note, multiple pixels share the signal originating from a sub-resolution emitter with varying amplitudes but identical lifetime signature. Grouping of pixels originating from the same particle therefore should enhance the phasor coordinate calculation (Fig. 1 *E-F*). When we applied this pixel binning strategy to the simulated data, we noticed a greatly reduced spread and a highly concentrated particle datapoint population in the phasor plot, now clearly centered on the 4 ns semicircle location. (Fig. 1 *G*). Subsequent color-coding of the simulated image for the particles that are detected with a sum of ≥300 photons showed that almost all particles were included into the analysis (Fig. 1 *H*). In summary, grouped single-particle phasor analysis helps accurately determine phasors for dim emitters, such as sub-resolution particles. Additionally, 300 photons per pixel binned particle is a recommended minimum total pixel count target for reliable phasor lifetime determination.

### Particle-based Phasor-FLIM resolves individual DNA molecule lifetimes

Next, we investigated whether particle-based phasor-FLIM could successfully allow determining the fluorescence lifetime of single molecules. Therefore, we imaged a model system containing PEG-Biotin immobilized DNA molecules labeled either with only a FRET donor (Fig. 2 *A-D*) or with both a FRET donor and acceptor spaced at a distance of 11 nucleotides as reported before (45, 46) (Fig. 2 *E-I*). All particles were identified, and particle phasor positions were analyzed in the phasor plot for the average lifetime. A donor only lifetime (𝜏_𝐷_) of 3.92 ns was found with a background (BG) contribution of 1.2% resulting in the measured phasor position (DO). Autofluorescence of the particles itself, here called BG, dragged an otherwise perfect monoexponential measurement inward of the semi-circle due to the multiexponential behavior of the autofluorescence. The BG position was determined (see Materials and Methods) and was used to Measure the pure lifetime of the donor on the semi-circle using fractionality (Fig. 2 *C-D*). With this no-FRET control measured we continued with a double labeled (donor-acceptor) dsDNA system, in which donor and acceptor labels are separated by 11 nucleotides (Fig. 2 *E*). These double labeled single-molecules were imaged using pulsed interleaved excitation (PIE), whereby the donor laser was pulsed at the beginning and our acceptor in the middle of our detection window in between pulses. This way quasi synchronous measurements of both channels were acquired without having the drawback of donor emission contaminating the acceptor channel. Colocalization was confirmed in a subset of single molecules using both donor and acceptor channel photon intensity information. Using a 5-pixel tolerance to identify double labeled particles 133 single molecule dsDNA particles were found (Fig. 2 *F*). From the colocalized particles, the donor signal upon donor excitation was used to calculate all phasor positions per particle using particle pixel binning. From the average position of the single (donor) labeled single dsDNA molecules (DO) and the previously measured BG coordinates (Fig. 2 *B*), a FRET trajectory was formed describing all possible FRET states ranging from 0% (phasor position equals the DO position) to 100% (phasor position equals the only remaining signal of BG; all donor signal is quenched) (Fig. 2 *G*). Finally, the FRET histogram resulting from projection of phasor positions onto the FRET trajectory showed an average value of 56% (Fig. 2 *H, I* and Fig. S1 *G*). Taking into account a Förster radius of 50.3 Å for Alexa 488 and ATTO 647N, the estimated donor-acceptor distance from our FRET measurement was determined to be 39 Å.

**Figure 2.**
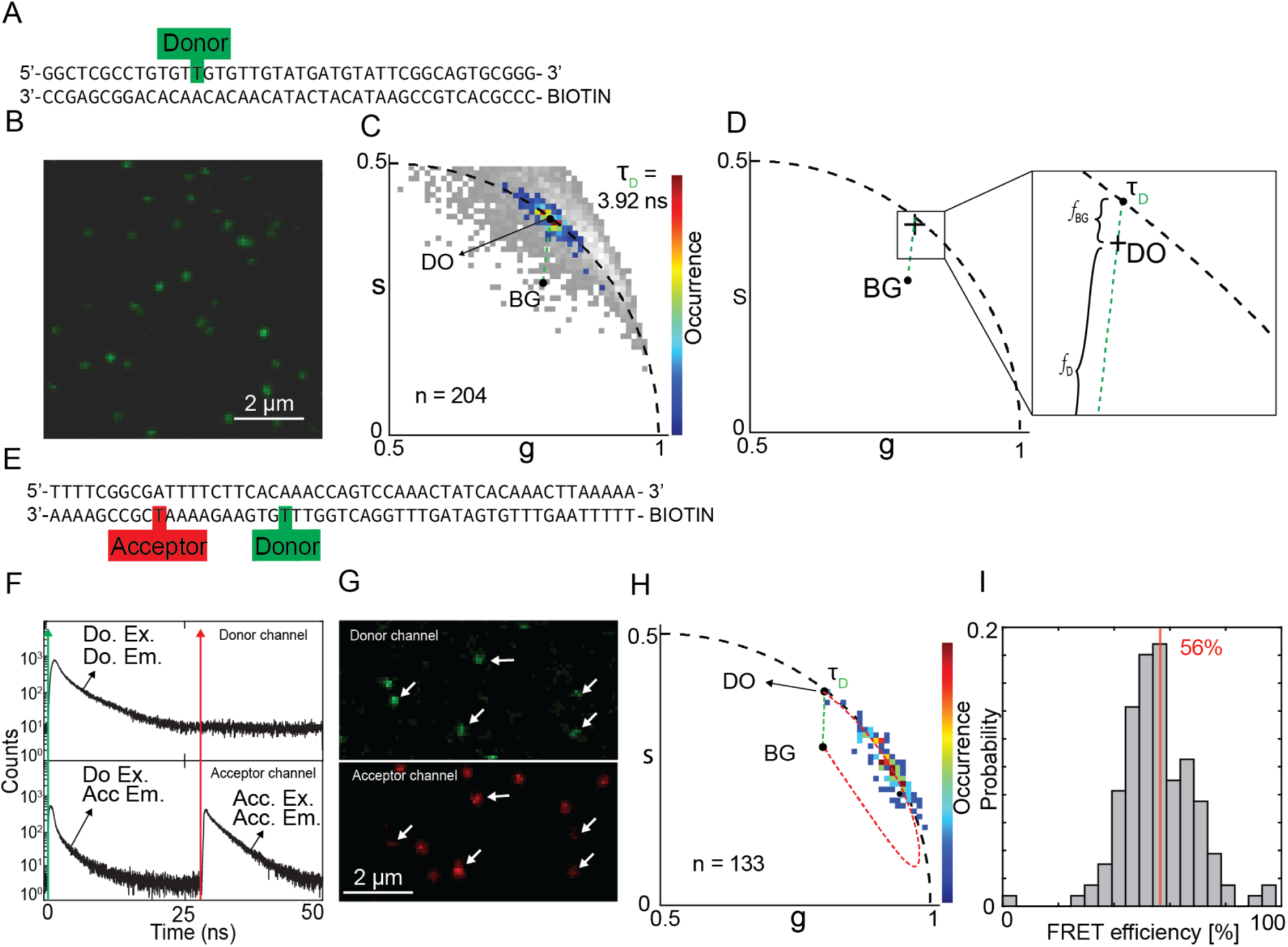
-Determination of FRET labeled dsDNA FRET efficiency: (A) dsDNA strand with a single donor label on the sense strand. (B) microscopy image using 485 nm excitation revealing single particles. (C) Pixel-binned particle phasor analysis on the donor-labeled dsDNA reveals its photon weighted center of mass (𝐷) revealing the donor lifetime of 3.92 ns (𝜏_𝐷_) upon extension of the 𝐵 − 𝐷 fraction line to reveal its pure contributing species. Pixel based phasor with a 10 photon/pixel threshold shown in gray. (D) Illustration of the contributing species to the phasor (𝐷) location (𝐵 and 𝜏_𝐷_) of the donor only labeled dsDNA. (E) dsDNA with on the antisense strand both donor and acceptor present and a biotin tag on the 5’ end used for immobilization on the glass. (F) Pulsed Interleaved Excitation (PIE) measurement of FRET labeled dsDNA used for colocalization. (G) Intensity image of double labeled dsDNA showing a single region of interest for both donor and acceptor channel as determined by colocalization of both donor and acceptor channels for donor and acceptor excitation respectively. White arrows indicate colocalized particles. (H) Phasor plot of imaged FRET labeled dsDNA strands showing the autofluorescence phasor (𝐵), the donor only phasor 𝐷, The quenching trajectory (red dashes) connecting the 𝐵 phasor with the 𝐷 phasor (see Eq. 22 & 23). The fraction line between 𝐵 en 𝐷 (green dashes). (I) FRET efficiency histogram derived from the FRET trajectory. Vertical red line is the average value.

### Performance comparison of single FP labeled integrase

With our positive FRET control in place, we connect to the open research field of HIV-1 integrase multimerization, previously conducted in our lab. Earlier, the FRET readout of recombinant HIV-1 particles containing fluorescently labelled IN (mTFP1-mVenus FRET pair) was investigated with intensity based acceptor photobleaching methods throughout the replication cycle (36–38). To extend this study using out phasor-FLIM particle approach we explored what integrase (IN) labels behave optimally for further application in such FRET measurements involving donor and acceptor labeled IN monomers.

Since an HIV-1 particle is a highly condensed environment and environment properties affect the fluorescence lifetime and photon output, we first needed to determine the optimal FRET pair in the HIV-1 particle context (47). This was achieved by measuring HIV-1 particles with a single type of labeled IN for several donor or acceptor labels. Firstly, we analyze the HIV-1 particles for their photon count and saw that especially particles containing IN-mTurquoise2 yield more 300+ photon particles as the frequency distribution peaks less in the low photon range (<100 counts). while mNeongreen does not contain many high photon count particles as the frequency distribution peaks at 30-40 photons (Fig. 3 *A*). For acceptor candidates the photon distributions are highly similar while IN-mCherry containing particles do provide the most high-photon shifted frequency histogram peak (Fig. 3 *E*).

**Figure 3:**
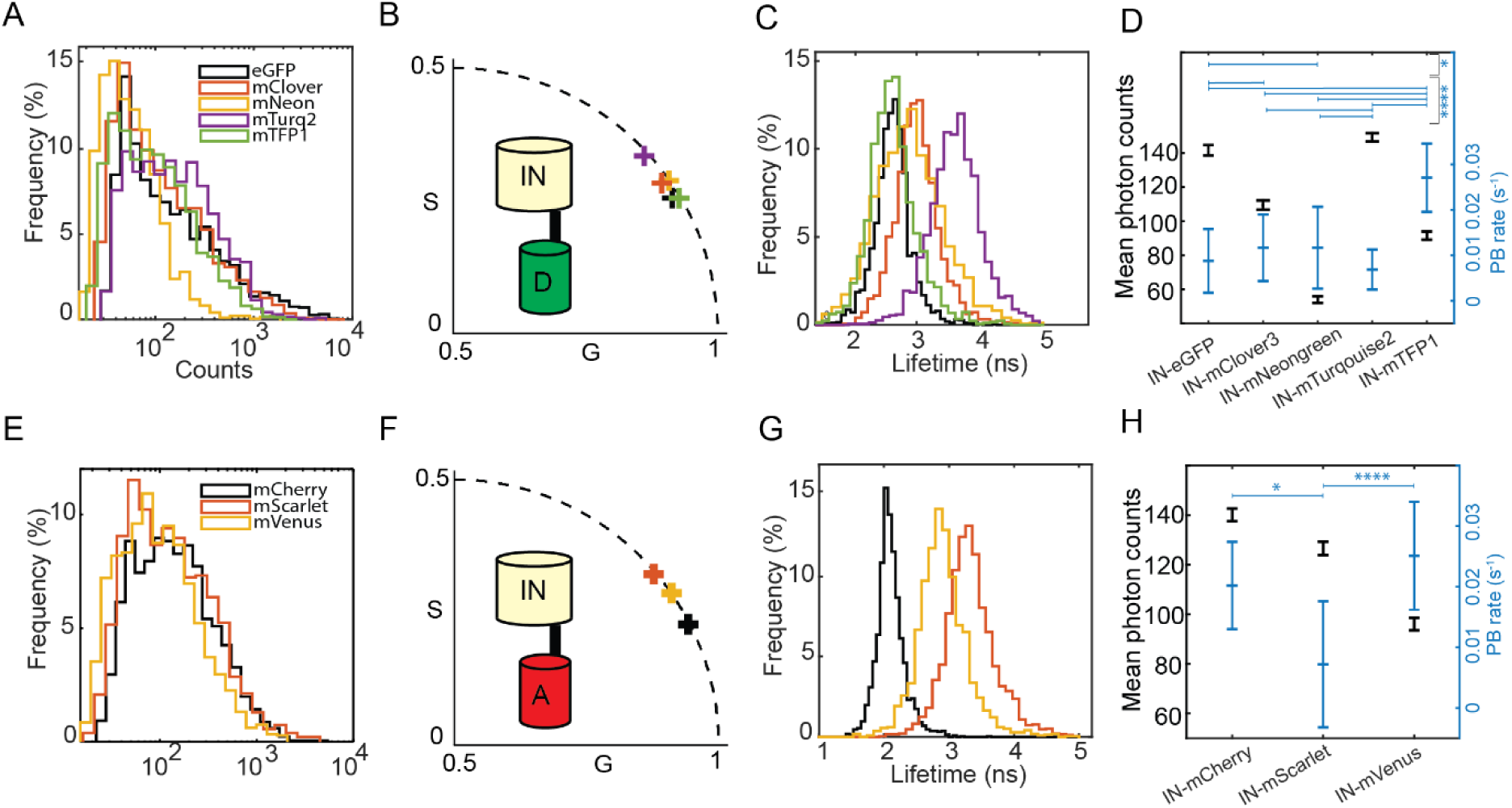
Lifetime, photon count and photobleaching study of single-FP HIV-IN labeled virions on glass. (A,E) Counts distribution of donor (A) and acceptor (E) labeled HIV-IN-FP virions. (B,F) Photon weighted centroid position of the phasor clouds of all respective donor and acceptor HIV-IN-FP virions. (C,G) Mono-exponential phase lifetime distribution of the donor and acceptor HIV-IN-FP virions using a particle threshold of 10 counts to evaluate broadness of lifetime distribution. (D,H) Geometric mean photon counts normalized for excitation efficiency and the associated mean photobleaching rate of all donor and acceptor HIV-IN-FP particles. All geometric mean photon counts are significantly different from each other (p<0.05), significance levels for photobleaching rates are determined by a one-way ANOVA non-paired non-parametric Kruskal-Wallis test with multiple comparison (* = p<0.05, **** = p<0.001).

Following, we verified that all IN-FP candidates, especially the donors, exhibit a mono exponential lifetime, which would enable us to use semi-circle derived lifetime values (phase or modulation lifetime would work) for evaluation. The plotted centroid positions of all measured particles in the phasor plot indicate that there is indeed a dominant mono exponential behavior (Fig. 3 *B,F*). Plotting all semi-circle derived phase lifetimes and their respective lifetime spread in a lifetime histogram shows their peak values and variability. Histogram means for each fluorophore were: eGFP 2.66±0.31 ns (n = 2675), mTFP1 2.62±0.41 ns (n= 1769), mNeongreen 2.90±0.60 ns (n= 1972), mClover3 3.01±0.39 ns (n= 2219), mTurquoise2 3.68±0.45 ns (n= 2375), mCherry 2.10±0.22 ns (n= 2164), mScarlet 3.34±0.4 ns (n = 2309) and mVenus 2.71±0.39 ns (n = 2125). With a variability of more than 0.5 ns, IN-mNeongreen particles are most unfavorable (Fig. 3 *C,G*). Measured photobleaching rates of the fluorescent proteins indicated a high rate for mTFP1 (0.0280 ± 0.0075, n = 29) while other donor candidates have a more favorable photobleaching rate: eGFP (0.0097 ± 0.0072, n = 143), mTurquoise2 (0.0078 ± 0.0044, n = 265), mClover3 (0.0127 ± 0.0073, n = 638), mNeongreen (0.0126 ± 0.0090, n = 112)(Fig. 3 *D*). When comparing the geometric mean photon counts all particles were significantly different from each other (p<0.001) and again eGFP (144.7, n = 2675) and mTurquoise2 (148.1, n = 2375) were best performing while mNeongreen (56.5, n = 1972), mClover3 (111.5, n = 2219) and mTFP1 (94.2, n = 1769) were significantly lower (all p<0.001)(Fig. 3 *D*). Regarding the measured acceptor photobleaching rates, mCherry (0.0202±0.0072, n = 15) and mVenus (0.0251 ± 0.0089, n = 17) exhibited a significantly larger photobleaching rate compared to mScarlet (0.0072 ± 0.0104, n = 14) (p<0.05 and p<0.001 respectively)

(Fig. 3 *H*). Mean photon counts were again all significantly different (p<0.001) and mCherry (118.7, n = 2164) and mScarlet (110.5, n = 2309) outperformed mVenus (90.1, n = 2125) (Fig. 3 *H*). Additionally, we point to the slight multiexponential character of these single labeled viral particles given their position inside the phasor semicircle due to autofluorescence contributions. Based on these results, we selected eGFP and mTurquoise2 as they appear to be favorable donor candidates that can be paired with acceptors from our assay that also score significantly. Due to its favorable photostability, mScarlet was shown to be most promising acceptor compared to mCherry and mVenus. In summary, we have identified suitable FRET donors and acceptors attached to integrase in the HIV-1 viral particles. The mTurquoise2-mVenus and eGFP-mScarlet pairs were selected for further single-particle FRET experiments.

### Graphical phasor analysis reveals integrase multimers in HIV-1 viral particles

IN is known to adopt different oligomeric states depending on the step in the replication cycle. Methods to quantify this oligomeric state of IN at the level of single viral particles and viral complexes are highly desired by virologists and measuring FRET with acceptor photobleaching intensity-based FRET has been conducted before. There, the dynamic range of these intensity-based FRET quantifications was unavoidably lowered by the presence of (variable amounts of) FRET donor-only labeled complexes. Using phasor-FLIM in the present work, we sought to further disentangle the actual FRET species from any non-FRET species and background.

Now, we employ the particle-based phasor-FLIM methodology on the FRET measurements for our previously selected fluorescent FRET pairs. These pairs each cover other sections of the spectrum, exhibit a different Förster distance (58.3 Å for mTurquoise2-mVenus and 56.7 Å for eGFP-mScarlet from calculations assuming κ² to be 2/3 and refractive index n of 1.33 with the donor quantum yields being 0.93 and 0.60 respectively) and encompass a FRET donor with substantially different lifetimes. Ultimately, regardless of the fluorescent labels on IN, we wanted to qualitatively evaluate the FRET state of IN complexes in the assembled virus particles to gain further insight into IN multimerization. Assuming a system where only monomeric IN and dimeric IN is present, in any combination possible (A, D, D:D, D:A, A:D, A:A), in a matured HIV-1 particle, there will be a passive donor intensity fraction between 50 % (all IN complexes are dimers, the donor-only signal intensity fraction constitutes 50% of the total donor signal intensity) and 100 % (all IN complexes are monomers, no FRET occurs). It is only when higher order multimers such as tetramers are formed, leading to a range of different combination between D and A (e.g., DDDA, DDAD, DADD, ADDD, ADAD, DADA), that the fraction of passive donors in IN complexes is further lowered, simultaneously increasing the overall FRET signal of a single HIV-1 particle. Since at the particle level both FRET and the passive donor fraction are unknown, no direct FRET readout can be provided. However, matching the particle phasor data with the possible FRET trajectories constructed by a range of passive donor contributions, should still give insights into the multimer composition. Practically, as shown in Fig. 4 *A* and *B,* we recorded FLIM data of FRET donor/acceptor double labeled immobilized viral particles at photon counts compatible with single-particle phasor-FLIM and performed particle detection on the resulting FRET donor images. FRET trajectories are constructed using a background reference (see Materials and Methods) and the HIV-1 particle measurements with only the donor label (HIV-1 containing only IN-mTurquoise2 or IN-eGFP, see Fig. *S10*). Again, the signal intensity contribution of BG in the measured donor signal draws the phasor position away from the semi-circle and serves as starting position for the FRET quenching line (as is previously illustrated in Fig. 2 *D*). For particles with IN-mTurquoise2 and IN-mVenus BG had a 7.5% contribution and for IN-eGFP/IN-mScarlet particles 10.5%, as determined graphically from the phasor plot. For these particles we next determined the donor signal phasor position per particle which are pooled into the 2D phasor histogram. The resulting FRET histograms derived from the 0%, 10% and 20% passive donor contribution FRET trajectories point towards a 12 and 32% FRET efficiency as average FRET value for the middle case with 10% passive donors (Fig. 4 *C, D*). Pixel based phasors of the same images is generated illustrating the difficulty of a graphical analysis method utilizing low photon count pixels (Fig. S11).

**Figure 4:**
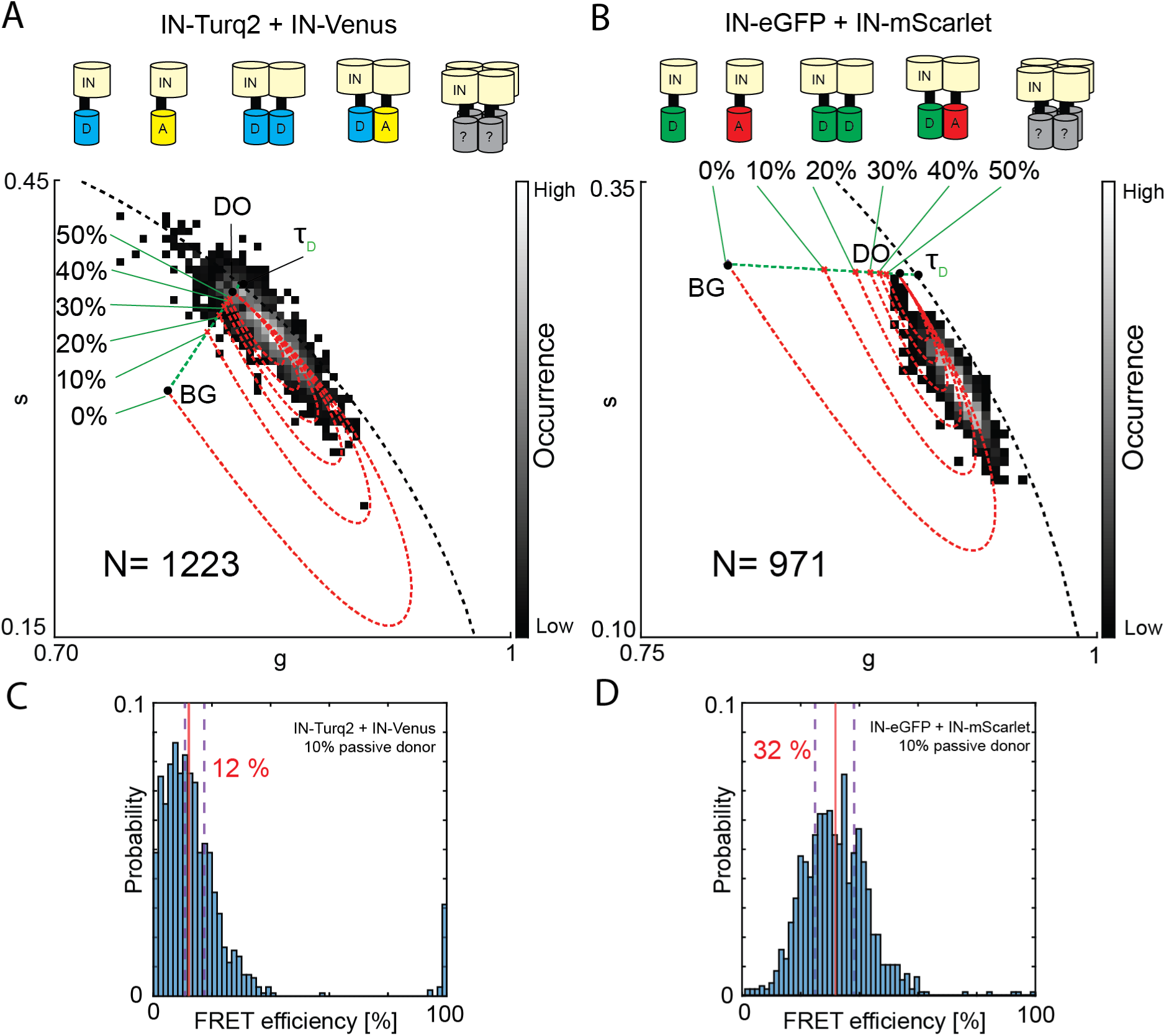
FRET trajectory analysis of HIV-1 particles containing donor labeled IN and acceptor labeled IN using particle-based pixel binning. (A) Phasor analysis of viral HIV-1 particles containing both donor and acceptor labeled IN (IN-Turquoise2+IN-mVenus) forming an unknown mix of multimers including monomers and dimers and potentially higher order multimers. The donor only phasor (DO) (g: 0.82, s:0.37 and background phasor (BG) (g: 0.78, s:0.31) determine the starting point and end of the FRET trajectory. Variations of the donor only signal contribution in the analyzed particles result in a variety of FRET trajectories and FRET trajectory end points (red dotted line). (B) Phasor analysis of viral HIV-1 particles containing both donor and acceptor labeled IN (IN-eGFP+IN-mScarlet) forming an unknown mix of multimers including monomers and dimers and potentially higher order multimers. The donor only phasor (DO) (g: 0.90, s:0.29) and background phasor (BG) (g: 0.79, s:0.31) determine the starting point and end of the FRET trajectory. Variations of the donor only signal contribution in the analyzed particles result in a variety of FRET trajectories and FRET trajectory end points (red dotted line). In both (A) and (B) the pure mono-exponential lifetime of the donor (𝜏_𝐷_) and BG determine by intensity fraction the position of DO on their connecting fraction line (green dotted line). (C) and (D) are FRET histograms extracted from the 0 %,10 % and 20 % passive donor contribution FRET trajectories of which the 0% and 20% histogram averages are indicated by purple dashed lines and the 10% passive donors histogram (shown) average are depicted as a vertical red line with its value.

## Discussion

In this paper, we demonstrate the usage of time domain phasor FLIM in combination with grouped pixel analysis for sub-resolution single particles. We applied this method on FRET labelled DNAs and on viral HIV-1 particles that contain single (donor) or double (donor and acceptor) labelled IN, which show energy transfer upon multimerization as demonstrated before using intensity based approaches (38, 44). First, we showed the importance of grouping pixels to obtain a higher quality phasor determination, optimizing the analysis to utilize all available photons of a particle. In a perfect setting (no background or shot noise) our simulations showed that 300 photons is the lower limit for a phasor determination within a 5% error range of the true lifetime of a single component. Therefore, when measuring with considerable background or working with complex multi-component samples, it is advised to aim for a higher photon count per object. The required photon budget is the weak point of every FLIM-FRET analysis compared with intensity based FRET approaches but is continuously improved with newer FLIM methodologies allowing more and more photons per pulse and a consequently faster imaging or more efficient detection (48). At the same time, using FLIM with PIE allows the robust combination of intensity based and FLIM quasi-simultaneously as is the case with smFRET burst measurements using multiparameter fluorescence detection with PIE (46, 49). Additionally PIE allows for multiplexing of (FRET) probes and provides flexibility in experimental design, such as three or even four color immobilized FRET (50) or using dark acceptor probes (51).

Currently, single object FLIM analysis is used by Qian et al. (35) for viral particle FLIM analysis and now further explored here with complex FRET behavior. We envision that this technique could be seen further applied in small organelles, protein complexes, immobilized single molecules or any system that provides homogeneous signal or signals that originate from a sub-resolution entity (and are therefore measured as average, potentially spread over multi pixels). Using object pixel binning allows for more precise analysis as one can oversample the image and later bin the pixels of desired structures. Up until now, images for single particle/molecule analysis are relatively under-sampled to obtain higher photon counts per pixel, voiding the image resolution.

As a remark, we measured our FLIM data using a 20MHz pulse rate to combine both acceptor and donor pulsing in the 50ns pulse window. When PIE is not required the used pulsing frequency can be changed to optimize usage of the phasor space for graphical purposes. A formula was provided by Clegg et al in 2005 for frequency domain phasor analysis (39):

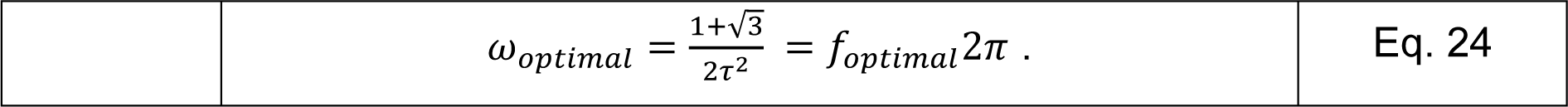

However, for time domain phasor applications the main improvement on the resolution of calculated s and g values will arise from a lowered TCSPC bin size.

Furthermore, the pixel-grouped FLIM approach was tested on labeled DNA structures, with two spaced fluorescent dyes set 11 nucleotides apart, to evaluate the detection and quantification of FRET by using the graphical phasor approach to FLIM. The obtained value of 39 Å is in close agreement with the literature, which described a value of 40 Å, determined by FRET restraint positioning and screening (FPS) simulations (46). Since a broad cross-lab study on dye-labeled DNAs pointed out their robustness, we deem our measurements as a validation of the analysis performance (45). We confirmed that the object binning strategy together with the phasor plot analysis allowed for determination of single molecule FRET via their lifetime signatures. This DNA sample provided a double labeled single molecule structure exhibiting a FRET signal from a known donor-acceptor stoichiometry, not complicating the measurement with a donor only fraction.

Following, we analyzed the performance of several fluorescent proteins, as it is difficult to predict the effect of IN coupling and the dense viral particle environment on the fluorescence brightness, lifetime and even maturation.

The donor fluorescent lifetimes were comparable to the known mono-exponential lifetimes found in literature for these FPs except mTFP1 which was 0.6 ns lower compared to the literature value of 3.2 ns (Table 1). The mCherry lifetime was also found to be inconsistent with literature, which could not be explained by the shifted ratios of the two exponential factors normally involved in a mCherry decay. Wu et al. described a two component lifetime for mCherry of 0.9 ns and 1.9 ns which are both lower than the measured 2.1 ns (61). mVenus and mScarlet on the other hand both measured 0.4 ns and 0.5 ns, which was above the normal literature values. In general, longer lifetimes are preferred for FRET because of higher dynamic range given a limited time resolution on the hardware side (53)(it is easier to measure a 2 ns difference compared to a 0.5 ns difference to determine a 50% FRET efficiency in a donor with a respective lifetime of 4 and 1 ns).

**Table 1:** Overview of measured single label IN in HIV-1 viral particles.

| FP | Literature lifetime | Measured lifetime | Measured photobleaching rate | Maturation (min) | Literature brightness | Measured geomean photon count | References |
| --- | --- | --- | --- | --- | --- | --- | --- |
| EGFP | 2.6 | 2.61 ± 0.31 | 0.0097 ± 0.0073 | 25 | 33.54 | 144.7 | (52)(53) |
| mClover3 | 3.2 | 3.01 ± 0.39 | 0.0127 ± 0.0073 | 43.5 | 85.02 | 111.5 | (53)(54) |
| mNeongreen | 3.1 | 2.90 ± 0.60 | 0.0126 ± 0.0090 | 10 | 92.8 | 56.5 | (54) |
| mTurquoise2 | 4.0 | 3.68 ± 0.45 | 0.0078 ± 0.0044 | 33.5 | 27.9 | 148.1 | (55) |
| mTFP1 | 3.2 | 2.62 ± 0.41 | 0.0280 ± 0.0075 | 76.5, | 54.4 | 94.2 | (56) |
| Cherry | 1.49* | 2.10 ± 0.22 | 0.0202 ± 0.0072 | 15 | 20 of 15.6 | 118.7 | (57)(58) |
| Scarlet | 3.86 | 3.34 ± 0.41 | 0.0072 ± 0.0104 | 174 | 71 | 110.5 | (54)(59) |
| Venus | 3.1 | 2.71 ± 0.39 | 0.0251 ± 0.0089 | 17.6 | 66.56 | 90.1 | (60) |

\*Averaged 2 component lifetime

Further, we expected a photostability double as good for mNeongreen compared to mClover3 (62). However, we recorded a similar photostability, which could be hinting to a photophysical problem for mClover3 or a gained benefit for mNeongreen in the HIV-1 particle environment. Looking at the particle photon counts, mClover3 and mNeongreen both underperformed while having the highest literature described brightness, further strengthening our belief that the emission of mClover3 is hampered in the viral particles. mTurquoise2 and mCherry performed well considering a generally low brightness. These constatations lead us to believe that fluorescent behaviors of IN-labeled proteins in HIV-1 are difficult to predict. We note that for some FPs of similar spectra identical filters were used and that the slightly different treatment of the filter of the emission spectrum could have introduced a small bias into our data.

Both viral particles types used in our final FRET experiment with HIV-1 particles, the resulting phasor cloud was found to be predominantly shifted towards a FRET value which is higher than any scenario in that matches 50% of the signal consisting of passive (non-FRET) donors, as would be the case with monomer-containing particles. Even for particles yielding a combination of monomers or dimers, the FRET readout cannot be explained as it would still be close to 50% passive donor signal. Therefore, in samples with equal numbers of IN-donor and IN-acceptor molecules this can only be explained by the presence of higher order multimers which reduce overall pure donor species since more are involved in FRET active oligomer.

While this is an honest and qualitative estimate, intensity-based FRET measurements would present the crude average of the active (FRET) and passive (no FRET) donors present in each viral particle. However, using the graphical analysis of the phasor plot remains challenging for a variety of reasons. Firstly, there remains a large error on the precise fraction of passive donors present in the particles, together with between-particle variation of that number. Additionally, every multimer formed in the imaged particles has its own FRET efficiency since it consists of a variable content of unlabeled and labeled (donor or acceptor) resulting in an extra variability in FRET readouts. Lastly, the composition of unlabeled, donor labeled and acceptor labeled cannot be verified, adding to difficulty of data interpretation. Overall, we can assume that measuring close to or more than 1000 particles provides a strong averaged readout with valuable information on the FRET state and therefore insight in the multimeric state of HIV-IN in assembled virus particles. One possible avenue of further investigation is redesigning particles with variable D-A ratios, variable total amounts of labels per particle, or non-FP labeling strategies altogether. Additionally, because the number of fluorescent proteins per particle is limited, the photon output is low and contributions of background and shot noise are considerable. We clearly show that taking the fraction of passive donors into account is essential. This factor is often overlooked because estimating the donor only fraction is difficult. Using the phasor analysis, we acknowledged the presence of passive donors in our viral particles, which will in any case, lower the apparent FRET signal measured. Given a more precise technique to determine the number of passive donors, researchers would be able to pinpoint which FRET trajectory is suitable and derive more absolute FRET efficiency values. At the same time, determining the number of passive donors present in a mixture of FRET displaying IN multimers remains a variable that can change particle to particle and can therefore not be fully generalized.

Although we cannot pinpoint the exact oligomerization state of every IN complex per particle, we can give a founded ensemble evaluation of the in vitro oligomeric state and expect to improve further when photon yield of such experiments becomes more efficient, increasing our phasor data quality. By trying to deconvolute the passive donor contribution we increase out analysis sensitivity to FRET changes, since the significant contribution can overshadow it otherwise.

Finally, while phasor-FLIM is said to be fit free, quenching trajectories and fraction lines are equally fitted onto the plot. Quantitative analysis in phasor plots equally fits graphical elements onto the data forcing the analyst to assume a data model. However, this does not eliminate the reciprocity of pixel-phasor and high level of data transparency that phasor exhibits to its users. We point to the requirements of correct phasor analysis which are not always equally straight forward to implement. Looking at the pixel-based phasor cloud, establishing the graphical phasor analysis where we can evaluate the position of single particle mixtures is impossible given enlarged data spreading due to low photons per pixel. Also, when one would look at the 2D histogram intensities for a given pixel-based phasor, graphical analysis is biased for the larger amount of low intensity pixels, which would require a photon weighted approach, diminishing the potential for an easy-to-implement graphical analysis.

Having established a very probable FRET value for the measured dual color HIV-1 particles, we presented a methodology for future use in following the FRET value of particles in several steps of the viral replication cycle, or it could prove profitable in other particle-based FRET research.

## Supporting material description

One supplemental note and eleven supplemental figures are available.

Supplemental note 1 describes the theory of concentration fractions in the phasor plot. Figure S1 gives a broad overview of the phasor theory for time domain FLIM.

Figure S2 illustrates the workflow in PAM software for data analysis and particle-based phasor as used.

Figure S3 Schematic of the used custom-built time-resolved confocal Figure S4 Illustration of the photon simulation UI used in PAM

Figure S5 Illustration of FCS and TauFit on organic dyes for microscope calibrations and phasor reference.

Figure S6 IN-eGFP particle time-trace fitted for photobleaching rate

Figure S7 Simulation experiment illustrating concentration fractions in phasor

Figure S8 Spatial color-coded illustration of phase and modulation within the semi-circle

Figure S9. Impact of IRF on phasor location without referencing

Figure S10 Donor only phasor-plots for single labeled HIV-1 integrase with mTurquoise2 and eGFP

Figure S11 Pixel based phasor analysis of FRET HIV-1 particles

## Acknowledgments

Jens de Wachter is thanked for assistance with the DNA experiments. Dr. Waldemar Schrimpf is thanked for implementing phasor analysis in PAM and evaluating the manuscript. Namjoo Vanderveken is acknowledged for assistance in producing the viral particle preparations. Rik Nuyts, Carine Jacker and Débora Linares are gratefully acknowledged for assistance in the lab and essential logistics. Dr. Kris Janssen is thanked for writing the software for galvanometric mirror scanning. Irena Zurnic Bönisch is acknowledged for critically reading the manuscript.

## Author contributions

GSF, JH and QC designed, built and aligned the microscope. QC and JH performed simulations. NP imaged the DNA. IZ produced the viral particles. NP and QC imaged viral particles. QC, CQ and JH adapted the PAM software for the specific analyses. NP performed preliminary analyses. QC, NP and JH performed all analyses. QC and JH wrote the manuscript. All authors reviewed and edited the manuscript.

## References

1. Heaster, T.M., J.T. Sharick, A.A. Gillette, M.C. Skala, R. Datta, T.M. Heaster, J.T. Sharick, A.A. Gillette, and M.C. Skala. 2020. Fluorescence lifetime imaging microscopy : fundamentals and advances in instrumentation, analysis, and applications. 25.

2. Chen, Y.C., B.Q. Spring, and R.M. Clegg. 2012. Fluorescence lifetime imaging comes of age how to do it and how to interpret it. Methods Mol. Biol. 875:1–22.

3. Berezin, M.M.Y., and S. Achilefu. 2010. Fluorescence Lifetime Measurements and Biological Imaging. Chem. Rev. 110:2641–2684.

4. Verveer, P.J., A. Squire, and P.I.H. Bastiaens. 2000. Global analysis of fluorescence lifetime imaging microscopy data. Biophys. J. 78:2127–2137.

5. Verveer, P.J., and P.I.H. Bastiaens. 2003. Evaluation of global analysis algorithms for single frequency fluorescence lifetime imaging microscopy data. J. Microsc. 209:1–7.

6. Digman, M.A., V.R. Caiolfa, M. Zamai, and E. Gratton. 2008. The phasor approach to fluorescence lifetime imaging analysis. Biophys. J. 94:L14–L16.

7. Hinde, E., M.A. Digman, C. Welch, K.M. Hahn, and E. Gratton. 2012. Biosensor Förster resonance energy transfer detection by the phasor approach to fluorescence lifetime imaging microscopy. Microsc. Res. Tech. 75:271–281.

8. Lou, J., A. Solano, Z. Liang, and E. Hinde. 2021. Phasor Histone FLIM-FRET Microscopy Maps Nuclear-Wide Nanoscale Chromatin Architecture With Respect to Genetically Induced DNA Double-Strand Breaks. Front. Genet. 12:1– 9.

9. Celli, A., S. Sanchez, M. Behne, T. Hazlett, E. Gratton, and T. Mauro. 2010. The epidermal Ca2+ gradient: Measurement using the phasor representation of fluorescent lifetime imaging. Biophys. J. 98:911–921.

10. Levitt, J.A., S.P. Poland, N. Krstajic, K. Pfisterer, A. Erdogan, P.R. Barber, M. Parsons, R.K. Henderson, and S.M. Ameer-Beg. 2020. Quantitative real-time imaging of intracellular FRET biosensor dynamics using rapid multi-beam confocal FLIM. Sci. Rep. 10:1–9.

11. Liu, Z., D. Pouli, C.A. Alonzo, A. Varone, S. Karaliota, K.P. Quinn, K. Mönger, K.P. Karalis, and I. Georgakoudi. 2018. Mapping metabolic changes by noninvasive, multiparametric, high-resolution imaging using endogenous contrast. Sci. Adv. 4.

12. Davis, R.T., K. Blake, D. Ma, M.B.I. Gabra, G.A. Hernandez, A.T. Phung, Y. Yang, D. Maurer, A.E.Y.T. Lefebvre, H. Alshetaiwi, Z. Xiao, J. Liu, J.W. Locasale, M.A. Digman, E. Mjolsness, M. Kong, Z. Werb, and D.A. Lawson. 2020. Transcriptional diversity and bioenergetic shift in human breast cancer metastasis revealed by single-cell RNA sequencing. Nat. Cell Biol. 22:310–320.

13. Datta, R., A. Alfonso-Garciá, R. Cinco, and E. Gratton. 2015. Fluorescence lifetime imaging of endogenous biomarker of oxidative stress. Sci. Rep. 5:1–10.

14. Stringari, C., H. Wang, M. Geyfman, V. Crosignani, V. Kumar, J.S. Takahashi, B. Andersen, and E. Gratton. 2015. In Vivo Single-Cell Detection of Metabolic Oscillations in Stem Cells. Cell Rep. 10:1–7.

15. Wright, B.K., L.M. Andrews, M.R. Jones, C. Stringari, M.A. Digman, and E. Gratton. 2012. Phasor-flim analysis of NADH distribution and localization in the nucleus of live progenitor myoblast cells. Microsc. Res. Tech. 75:1717–1722.

16. Stringari, C., R.A. Edwards, K.T. Pate, M.L. Waterman, P.J. Donovan, and E. Gratton. 2012. Metabolic trajectory of cellular differentiation in small intestine by Phasor Fluorescence Lifetime Microscopy of NADH. Sci. Rep. 2.

17. Schrimpf, W., J. Jiang, Z. Ji, P. Hirschle, D.C. Lamb, O.M. Yaghi, and S. Wuttke. 2018. Chemical diversity in a metal-organic framework revealed by fluorescence lifetime imaging. Nat. Commun. 9.

18. Fereidouni, F., A.N. Bader, and H.C. Gerritsen. 2012. Spectral phasor analysis allows rapid and reliable unmixing of fluorescence microscopy spectral images. Opt. Express. 20:12729.

19. Scipioni, L., A. Rossetta, G. Tedeschi, and E. Gratton. 2021. Phasor S-FLIM: a new paradigm for fast and robust spectral fluorescence lifetime imaging. Nat. Methods. 18:542–550.

20. Ranjit, S., L. Lanzano, and E. Gratton. 2014. Mapping diffusion in a living cell via the phasor approach. Biophys. J. 107:2775–2785.

21. Scipioni, L., E. Gratton, A. Diaspro, and L. Lanzanò. 2016. Phasor Analysis of Local ICS Detects Heterogeneity in Size and Number of Intracellular Vesicles. Biophys. J. 111:619–629.

22. Chiu, C.L., and E. Gratton. 2013. Axial super resolution topography of focal adhesion by confocal microscopy. Microsc. Res. Tech. 76:1070–1078.

23. Verveer, P.J., A. Squire, and P.I.H. Bastiaens. 2000. Global Analysis of Fluorescence Lifetime Imaging Microscopy Data. 78:2127–2137.

24. Pelet, S., M.J.R. Previte, L.H. Laiho, and P.T.C. So. 2004. A Fast Global Fitting Algorithm for Fluorescence Lifetime Imaging Microscopy Based on Image Segmentation. Biophys. J. 87:2807–2817.

25. Rahim, M.K., J. Zhao, H. V. Patel, H.A. Lagouros, R. Kota, I. Fernandez, E. Gratton, and J.B. Haun. 2022. Phasor Analysis of Fluorescence Lifetime Enables Quantitative Multiplexed Molecular Imaging of Three Probes. Anal. Chem. 94:14185–14194.

26. Albertazzi, L., D. Arosio, L. Marchetti, F. Ricci, and F. Beltram. 2009. Quantitative FRET analysis with the E0GFP-mCherry fluorescent protein pair. Photochem. Photobiol. 85:287–297.

27. De Los Santos, C., C.W. Chang, M.A. Mycek, and R.A. Cardullo. 2015. FRAP, FLIM, and FRET: Detection and analysis of cellular dynamics on a molecular scale using fluorescence microscopy. Mol. Reprod. Dev. 82:587–604.

28. Maus, M., M. Cotlet, J. Hofkens, T. Gensch, F.C. De Schryver, J. Schaffer, and C.A.M. Seidel. 2001. An experimental comparison of the maximum likelihood estimation and nonlinear least-squares fluorescence lifetime analysis of single molecules. Anal. Chem. 73:2078–2086.

29. Cotlet, M., S. Masuo, G. Luo, J. Hofkens, M. Van Der Auweraer, J. Verhoeven, K. Müllen, S.X. Xiaoliang, and F. De Schryver. 2004. Probing conformational dynamics in single donor-acceptor synthetic molecules by means of photoinduced reversible electron transfer. Proc. Natl. Acad. Sci. U. S. A. 101:14343–14348.

30. Kuimova, M.K., S.W. Botchway, A.W. Parker, M. Balaz, H.A. Collins, H.L. Anderson, K. Suhling, and P.R. Ogilby. 2009. Imaging intracellular viscosity of a single cell during photoinduced cell death. Nat. Chem. 1:69–73.

31. Okabe, K., N. Inada, C. Gota, Y. Harada, T. Funatsu, and S. Uchiyama. 2012. Intracellular temperature mapping with a fluorescent polymeric thermometer and fluorescence lifetime imaging microscopy. Nat. Commun.

32. Agronskaia, A. V, L. Tertoolen, and H.C. Gerritsen. 2004. Fast fluorescence lifetime imaging of calcium in living cells. .

33. Goryashchenko, A.S., A.A. Pakhomov, A. V. Ryabova, I.D. Romanishkin, E.G. Maksimov, A.N. Orsa, O. V. Serova, A.A. Mozhaev, M.A. Maksimova, V.I. Martynov, A.G. Petrenko, and I.E. Deyev. 2021. FLIM-Based Intracellular and Extracellular pH Measurements Using Genetically Encoded pH Sensor. Biosensors. 11.

34. Lin, H.J., P. Herman, and J.R. Lakowicz. 2003. Fluorescence Lifetime-Resolved pH Imaging of Living Cells. Cytometry. A. 52:77.

35. Qian, C., A. Flemming, B. Müller, and D.C. Lamb. 2022. Dynamics of HIV-1 Gag Processing as Revealed by Fluorescence Lifetime Imaging Microscopy and Single Virus Tracking. Viruses. 14.

36. Desimmie, B.A., R. Schrijvers, J. Demeulemeester, D. Borrenberghs, C. Weydert, W. Thys, S. Vets, B. Van Remoortel, J. Hofkens, J. De Rijck, J. Hendrix, N. Bannert, R. Gijsbers, F. Christ, and Z. Debyser. 2013. LEDGINs inhibit late stage HIV-1 replication by modulating integrase multimerization in the virions. Retrovirology. 10:1–16.

37. 37. Borrenberghs, D., W. Thys, S. Rocha, J. Demeulemeester, C. Weydert, P. Dedecker, J. Hofkens, Z. Debyser, and J. Hendrix. 2014. HIV Virions as Nanoscopic Test Tubes for Probing Oligomerization of the Integrase Enzyme. .

38. Borrenberghs, D., L. Dirix, F. De Wit, S. Rocha, J. Blokken, S. De Houwer, R. Gijsbers, F. Christ, J. Hofkens, J. Hendrix, and Z. Debyser. 2016. Dynamic Oligomerization of Integrase Orchestrates HIV Nuclear Entry. Sci. Rep. 6:1–14.

39. Redford, G.I., and R.M. Clegg. 2005. Polar plot representation for frequency-domain analysis of fluorescence lifetimes. J. Fluoresc. 15:805–815.

40. Lakowicz, J.R., H. Szmacinski, K. Nowaczyk, K.W. Berndt, and M. Johnson. 1992. Fluorescence lifetime imaging. Anal. Biochem. 202:316–330.

41. Gadella, T.W.J., T.M. Jovin, and R.M. Clegg. 1993. Fluorescence lifetime imaging microscopy (FLIM): Spatial resolution of microstructures on the nanosecond time scale. Biophys. Chem. 48:221–239.

42. Schrimpf, W., A. Barth, J. Hendrix, and D.C. Lamb. 2018. PAM: A Framework for Integrated Analysis of Imaging, Single-Molecule, and Ensemble Fluorescence Data. Biophys. J. 114:1518–1528.

43. Albanese, A., D. Arosio, M. Terreni, and A. Cereseto. 2008. HIV-1 Pre-Integration Complexes Selectively Target Decondensed Chromatin in the Nuclear Periphery. PLoS One. 3:2413.

44. Borrenberghs, D., W. Thys, S. Rocha, J. Demeulemeester, C. Weydert, P. Dedecker, J. Hofkens, Z. Debyser, and J. Hendrix. 2014. HIV virions as nanoscopic test tubes for probing oligomerization of the integrase enzyme. ACS Nano. 8:3531–3545.

45. Hellenkamp, B., S. Schmid, O. Doroshenko, O. Opanasyuk, R. Kühnemuth, S. Rezaei Adariani, B. Ambrose, M. Aznauryan, A. Barth, V. Birkedal, M.E. Bowen, H. Chen, T. Cordes, T. Eilert, C. Fijen, C. Gebhardt, M. Götz, G. Gouridis, E. Gratton, T. Ha, P. Hao, C.A. Hanke, A. Hartmann, J. Hendrix, L.L. Hildebrandt, V. Hirschfeld, J. Hohlbein, B. Hua, C.G. Hübner, E. Kallis, A.N. Kapanidis, J.Y. Kim, G. Krainer, D.C. Lamb, N.K. Lee, E.A. Lemke, B. Levesque, M. Levitus, J.J. McCann, N. Naredi-Rainer, D. Nettels, T. Ngo, R. Qiu, N.C. Robb, C. Röcker, H. Sanabria, M. Schlierf, T. Schröder, B. Schuler, H. Seidel, L. Streit, J. Thurn, P. Tinnefeld, S. Tyagi, N. Vandenberk, A.M. Vera, K.R. Weninger, B. Wünsch, I.S. Yanez-Orozco, J. Michaelis, C.A.M. Seidel, T.D. Craggs, and T. Hugel. 2018. Precision and accuracy of single-molecule FRET measurements—a multi-laboratory benchmark study. Nat. Methods. 15:669–676.

46. Vandenberk, N., A. Barth, D. Borrenberghs, J. Hofkens, and J. Hendrix. 2018. Evaluation of Blue and Far-Red Dye Pairs in Single-Molecule Förster Resonance Energy Transfer Experiments. J. Phys. Chem. B. 122:4249–4266.

47. Toseland, C.P. 2013. Fluorescent labeling and modification of proteins. J. Chem. Biol. 6:85–95.

48. Alvarez, L.A.J., B. Widzgowski^1^, G. Ossato, B. Van Den Broek, K. Jalink, L. Kuschel, M.J. Roberti, and F. Hecht. 2019. SP8 FALCON: a novel concept in fluorescence lifetime imaging enabling video-rate confocal FLIM. Nat. Methods. 20:2–4.

49. Hendrix, J., and D.C. Lamb. 2013. Pulsed interleaved excitation: Principles and applications. 1st ed. Elsevier Inc.

50. Ratzke, C., B. Hellenkamp, and T. Hugel. 2014. Four-colour FRET reveals directionality in the Hsp90 multicomponent machinery. Nat. Commun. 2014 *51*. 5:1–9.

51. Laskaratou, D., G.S. Fernández, Q. Coucke, E. Fron, S. Rocha, J. Hofkens, J. Hendrix, and H. Mizuno. 2021. Quantification of FRET-induced angular displacement by monitoring sensitized acceptor anisotropy using a dim fluorescent donor. Nat. Commun. 12:1–12.

52. Cormack, B.P., R.H. Valdivia, and S. Falkow. 1996. FACS-optimized mutants of the green fluorescent protein (GFP). Gene. 173:33–38.

53. Martin, K.J., E.J. McGhee, J.P. Schwarz, M. Drysdale, S.M. Brachmann, V. Stucke, O.J. Sansom, and K.I. Anderson. 2018. Accepting from the best donor; analysis of long-lifetime donor fluorescent protein pairings to optimise dynamic FLIM-based FRET experiments. PLoS One. 13:1–25.

54. McCullock, T.W., D.M. MacLean, and P.J. Kammermeier. 2020. Comparing the performance of mScarlet-I, mRuby3, and mCherry as FRET acceptors for mNeonGreen. PLoS One. 15:1–22.

55. Goedhart, J., D. Von Stetten, M. Noirclerc-Savoye, M. Lelimousin, L. Joosen, M.A. Hink, L. Van Weeren, T.W.J. Gadella, and A. Royant. 2012. Structure-guided evolution of cyan fluorescent proteins towards a quantum yield of 93%. Nat. Commun. 3.

56. Guerra, P., L.A. Vuillemenot, B. Rae, V. Ladyhina, and A. Milias-Argeitis. 2022. Systematic in Vivo Characterization of Fluorescent Protein Maturation in Budding Yeast. ACS Synth. Biol. 11:1129–1141.

57. Mukherjee, S., P. Manna, S.T. Hung, F. Vietmeyer, P. Friis, A.E. Palmer, and R. Jimenez. 2022. Directed Evolution of a Bright Variant of mCherry: Suppression of Nonradiative Decay by Fluorescence Lifetime Selections. J. Phys. Chem. B. 126:4659–4668.

58. Shaner, N.C., R.E. Campbell, P.A. Steinbach, B.N.G. Giepmans, A.E. Palmer, and R.Y. Tsien. 2004. Improved monomeric red, orange and yellow fluorescent proteins derived from Discosoma sp. red fluorescent protein. Nat. Biotechnol. 22:1567–1572.

59. Bindels, D.S., L. Haarbosch, L. Van Weeren, M. Postma, K.E. Wiese, M. Mastop, S. Aumonier, G. Gotthard, A. Royant, M.A. Hink, and T.W.J. Gadella. 2016. MScarlet: A bright monomeric red fluorescent protein for cellular imaging. Nat. Methods. 14:53–56.

60. Kremers, G.J., J. Goedhart, E.B. Van Munster, and T.W.J. Gadella. 2006. Cyan and yellow super fluorescent proteins with improved brightness, protein folding, and FRET förster radius. Biochemistry. 45:6570–6580.

61. Wu, B., Y. Chen, and J.D. Müller. 2009. Fluorescence fluctuation spectroscopy of mCherry in living cells. Biophys. J. 96:2391–2404.

62. Shaner, N.C., G.G. Lambert, A. Chammas, Y. Ni, P.J. Cranfill, M.A. Baird, B.R. Sell, J.R. Allen, R.N. Day, M. Israelsson, M.W. Davidson, and J. Wang. 2013. A bright monomeric green fluorescent protein derived from Branchiostoma lanceolatum. Nat. Methods. 10:407–409.

