## Supplementary Information for "Particle-based phasor-FLIM-FRET resolves protein-protein interactions inside single viral particles"

[$] Shared authorship

* Corresponding author

Last updated: May 30th, 2023

### Contents:


- Supplemental note 1 describes the theory of concentration fractions in the phasor plot.
- Figure S1 gives a broad overview of the phasor theory for time domain FLIM.
- Figure S2 illustrates the workflow in PAM software for data analysis and particle-based phasor as used.
- Figure S3 provides schematic of the used custom-built time-resolved confocal
- Figure S4 provides an illustration of the photon simulation UI used in PAM
- Figure S5 FCS and TauFit on organic dyes for microscope calibration and phasor reference.
- Figure S6 IN-eGFP particle time-trace fitted for photobleaching rate
- Figure S7 simulation experiment illustrating concentration fractions in phasor
- Figure S8 spatial color-coded illustration of phase and modulation within the semi-circle
- Figure S9. impact of IRF on phasor location without referencing
- Figure S10 donor only phasor-plots for single labeled HIV-1 integrase with mTurquoise2 and eGFP
- Figure S11 pixel-based phasor analysis of FRET HIV-1 particles

#### Supplemental note 1

In the case an unquenched species with a mono-exponential decay co-exists in the same pixel with a quenched form, the fluorescence fraction of one species $f$ is not equal to the concentration fraction $f_C$ when looking at the fraction line connecting both phasors. The formula used to determine the concentration fraction in this case can be derived as following. A pure fluorescence species with mono-exponential decay can be approximated by the following equation, omitting a possible effect of the instrument response function (IRF).

| | | |
|---|---|---|
| | $I(t) = Aexpexp\left[\frac{-t}{\tau}\right]$ | Eq. S1 |

Here the intensity over time of the decay ($I(t)$) follows from an exponential with amplitude $A$ (proportional to the concentration), lifetime $\tau$ and time $t$.

For a mixture of two species with different lifetimes, this gives:

| | | |
|---|---|---|
| | $I(t) = A_1.exp\left[\frac{-t}{\tau_1}\right] + A_2.exp\left[\frac{-t}{\tau_2}\right]$ | Eq. S2 |

The concentration fraction of species 1 is simply given by:

| | | |
|---|---|---|
| | $f_{c,1} = \frac{A_1}{A_1 + A_2}$ | Eq. S3 |

The fraction of photons radiating from species 1, i.e., the intensity fraction of species 1, on the other hand, is given by:

| | | |
|---|---|---|
| [1] | $f_1 = \frac{A_1\tau_1}{A_1\tau_1 + A_2\tau_2},$ | Eq. S4 |

We now rearrange this equation as follows:

| | | |
|---|---|---|
| | $\frac{1}{f_1} - 1 = \frac{A_2\tau_2}{A_1\tau_1}$ | Eq. S5 |

Since the ratio of amplitudes is the same as the ratio of concentrations, the following is true:

| | | |
|---|---|---|
| | $\frac{\tau_1}{\tau_2}\left(\frac{1}{f_1} - 1\right) = \frac{A_2}{A_1} = \frac{f_{c,2}}{f_{c,1}} = \frac{1 - f_{c,1}}{f_{c,1}} = \frac{1}{f_{c,1}} - 1$ | Eq. S6 |

Hence, the concentration fraction of species 1 in the mixture can be calculated from the different lifetimes and their corresponding intensity fractions:

| | | |
|---|---|---|
| | $f_{c,1} = \left(\frac{\tau_1}{\tau_2}\left(\frac{1}{f_{1,intensity}} - 1\right) + 1\right)^{-1}$ | Eq. S7 |


[1] J.R. Lakowicz, Springer US, 2006.

#### Supplemental Figure 1

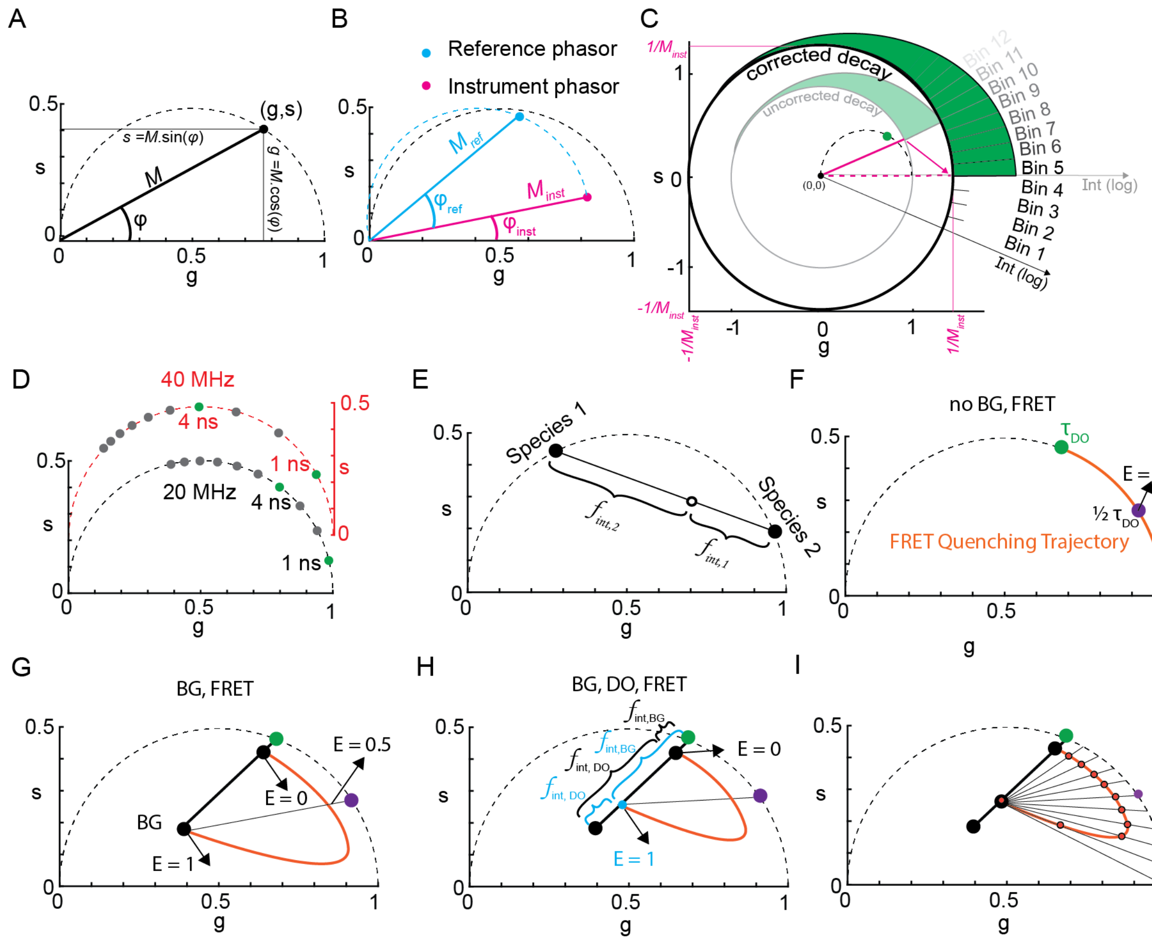


Figure S1: Overview figure of phasor theory for time-domain FLIM. (A) Every phasor, or phase vector, in the phasor space have two ways to be described. Firstly, the phase and modulation of the signal can be determined and used to plot a vector with a length $M$ and angle $\varphi$. Second, phasors can be described using cartesian coordinates $g$ and $s$ which represent the sine and cosine Fourier transform of the signal. Phasors on the semicircle represent mono-exponential lifetimes and within multi-exponential. A phasor which falls outside of the semicircle is the result of a non-exponential process. (B) Direct Fourier transform of a fluorescence decay results in a misplaced phasor. A phasor of a single exponential will not be found on the semicircle since the microscope is not a perfect machine. IRF and decay start vary in placement within the used TAC-range. When a reference sample with known mono-exponentiality and a known lifetime is measured the theoretical $M_{ref}$ and $\varphi_{ref}$ is calculated. These values most often do not land on the expected phasor location at the semi-circle due to convolution with the instrument response function. Using the expected $M$ and $\varphi$ of the known lifetime sample, the required change in the eventual phasor coordinate system is retrieved and can subsequently be used in later sample measurements. (C) By shifting and resizing the universal circle (centered around $(0,0)$ with a radius of 1) we make sure that any monoexponential will end up on the semicircle and that $(g_{inst}, s_{inst})$ is located at $(1,0)$, at a zero lifetime. The universal circle carries all TCSPC bins in the TAC range are spread over the circle circumference starting at $(1,0)$. Every position on the universal circle, representing a TAC-bin, is weighted by the photons that arrived in that bin. The average position of all photon-weighted universal circle positions determines the phasor position. Hence, oversizing and rotating the universal circle can counter the instrument contribution leading to correct representation of lifetime in and on the semicircle starting in $(1,0)$ at zero lifetime and nearing infinite lifetimes at $(0,0)$. The infinity lifetime at $(0,0)$ is achieved when a time domain measurement has no decay. This equal contribution of all bin-positions on the universal circle will average out at $(0,0)$. (D) Depending on the used TAC range, lifetimes are distributed accordingly for 20Mhz (black) and 40Mhz (red). (E) a combination of fluorescent species, a biexponential in the case of two species, will be located within the semi-circle and on the line connecting both species. Its position is the result of fractional contribution of photons from both species. (F) A FRET quenching line is used to represent all positions for E= 0 to 1 of a donor quenched by an acceptor. In case of no background ($BG$) and donor molecules not taking part in the FRET process ($DO$) the possible phasors follow the semicircle towards $(0,1)$. (G) When BG is convoluted within the measured decay, there is an increasing contribution of $BG$ as the quenched donor reduces its intensity fraction. At the point of fully quenched donor, the quenching line end at the only contributing species left, $BG$. (H) In case there are donor molecules not taking part in FRET (passive donors whose phasor location is $DO$) and there is $BG$ within the measured pixel decay the starting position, affected by BG, and the end position, affected by BG and DO, of the quenching line are adjusted forming a more compact trajectory. Fractional contributions of the situation at E=0 and E=1 are shown in black and blue respectively (I) Every point on the quenching line is determined from the fractionality between the phasor determined by $BG + DO$ and the quenched species phasor on the semicircle leading to rapidly increasing FRET efficiencies towards the end of the quenching line.

#### Supplemental Figure 2

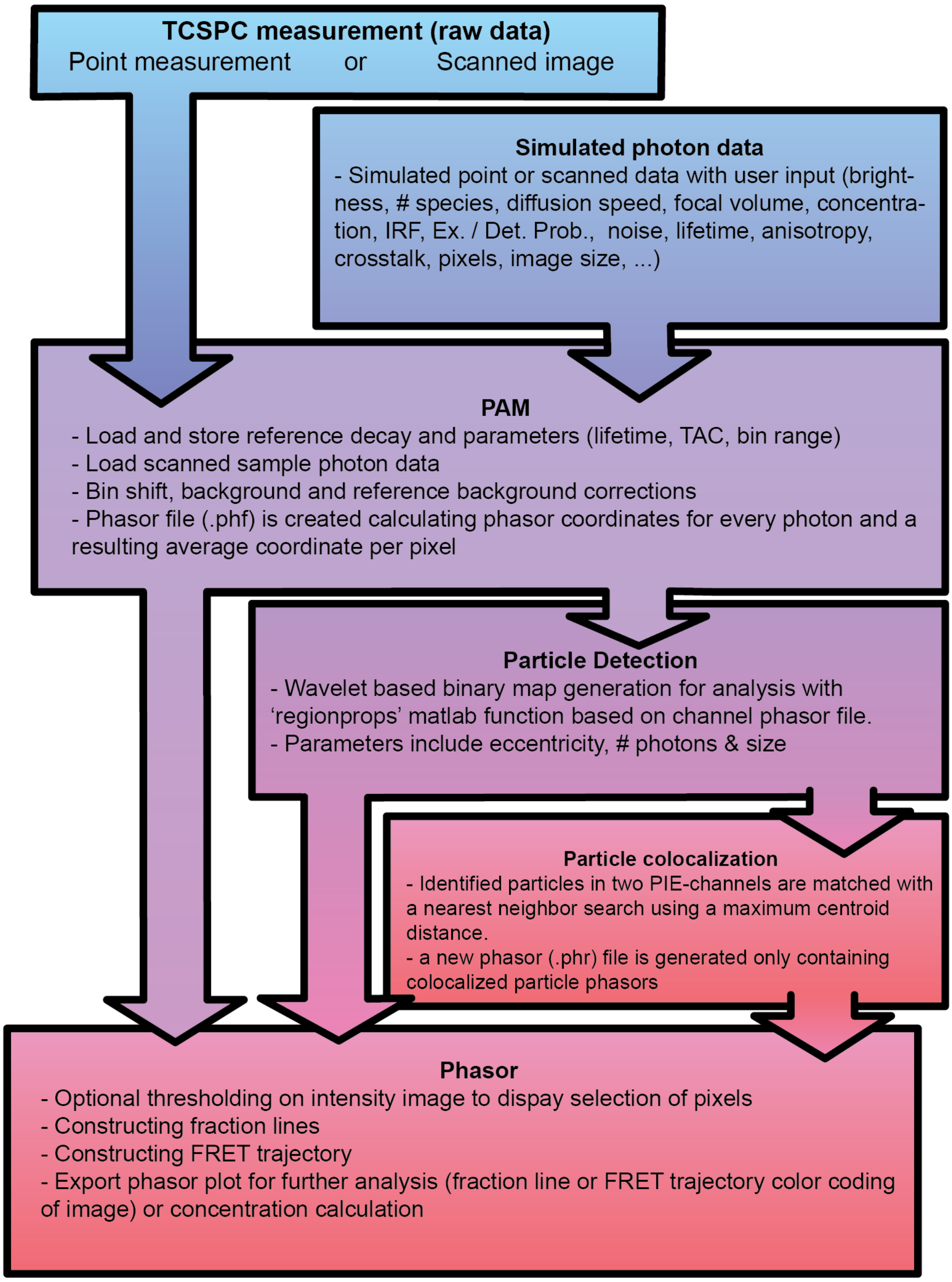


Figure S2: Flow-chart of the PAM software package showing the modules used in their respective order.

### Supplemental Figure 3

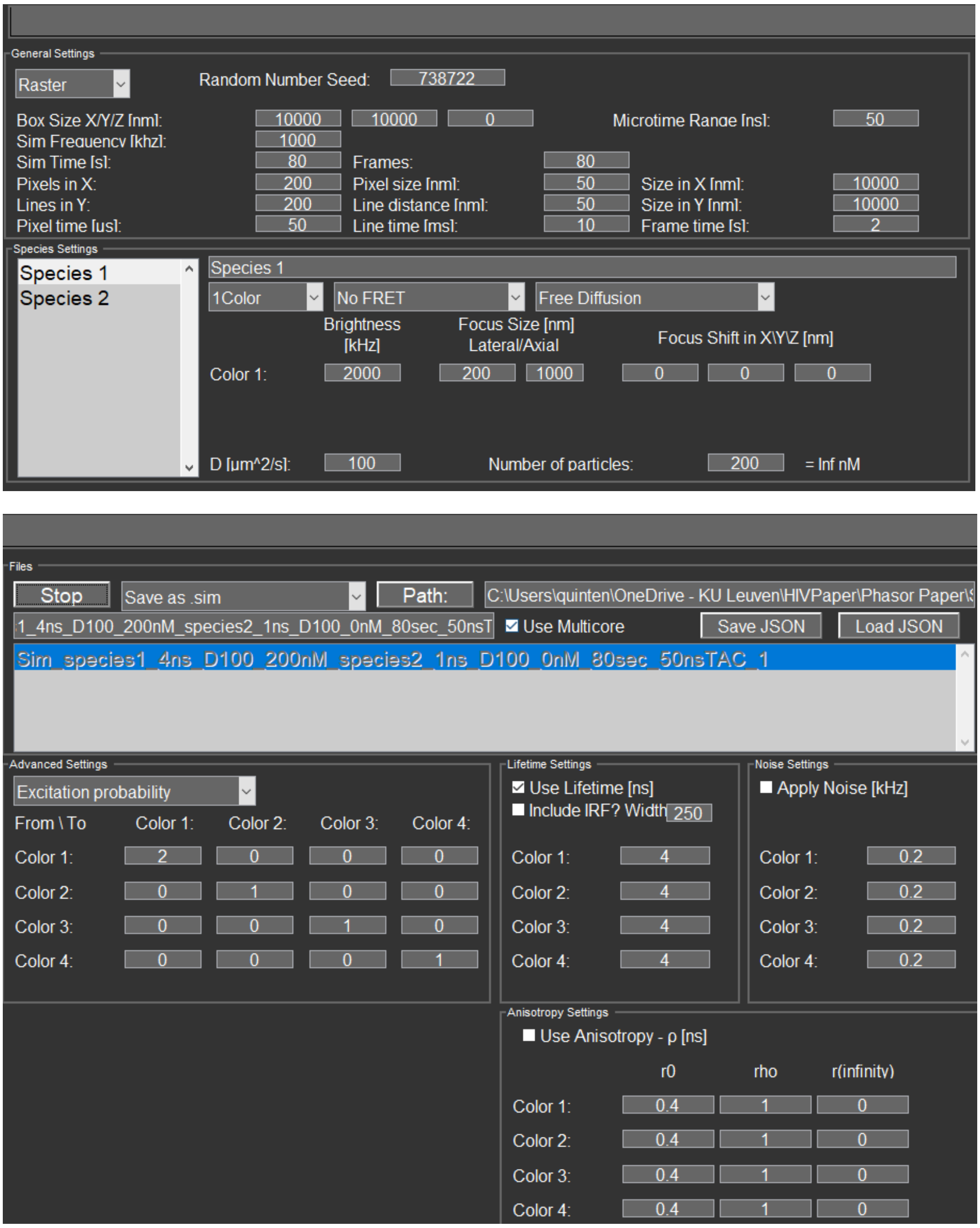


Figure S3: UI of the PAM simulation module. In the General setting, a simbox can be chosen in XYZ, scan settings, imaging duration and pulse frequency is set. In the species settings tab multiple species can be added, each with a desired brightness, diffusion speed and focus size. Additionally, the user chooses to include a one- or multicolor species, a static or dynamic FRET model and chooses from a list of diffusion types. Advanced settings allow to set the excitation probability, crosstalk and detection and bleaching probabilities. Finally, color-bound anisotropy lifetime and noise info can be given as input and an IRF can be convoluted into the simulated photon data.

#### Supplemental Figure 4

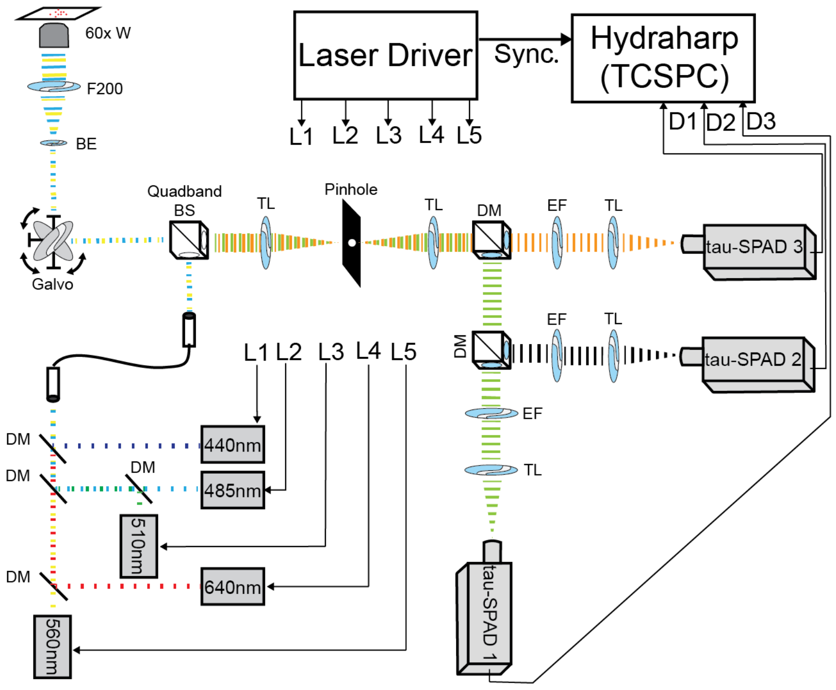


Figure S4: Schematic of a custom-built time resolved confocal setup. Abbreviations used: (W) water, (F200) lens with focus distance of 200mm, (BE) beam expander, (DM) dichroic mirror, (TL) tube lens, (EF) emission filter, (L) laser line, (D) detection line, (BS) beam splitter. All laser excitation light is shown up to the final combination mirror, from there we illustrated the 485 nm and 560 nm as used for an Alexa 488-Atto 647N FRET experiment with detection separated by one 560 nm long pass filter. In this case the second magnetic DM holder is empty, letting the fluorescence pass.

### Supplemental Figure 5

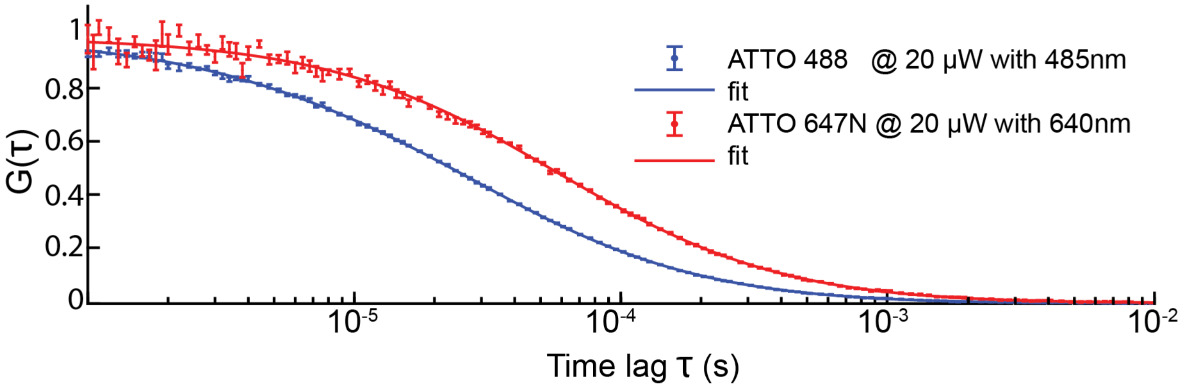


| File | Active | Counts [kHz] | Brightness [kHz] | N | F | G | D[$\mu m^2/s$] | F | G | $w_r$[µm] | F | G | $w_z$[µm] | F | G | tauT[µs] | F | G | Trip | F | G | y0 | F | G | Chi2 |
|---|---|---|---|---|---|---|---|---|---|---|---|---|---|---|---|---|---|---|---|---|---|---|---|---|---|
| atto488_ | ☑ | 32.0957 | 113.81 | 0.28201 | ☐ | ☐ | 373.379 | ☐ | ☐ | 0.20809 | ☐ | ☐ | 1.0794 | ☐ | ☐ | 3.9811 | ☐ | ☐ | 0.072399 | ☐ | ☐ | 0.0008834 | ☐ | ☐ | 0.8442 |
| atto647n | ☑ | 15.7911 | 34.2385 | 0.46121 | ☐ | ☐ | 341.4929 | ☐ | ☐ | 0.2836 | ☐ | ☐ | 1.3086 | ☐ | ☐ | 4.3437 | ☐ | ☐ | 0.014432 | ☐ | ☐ | 0.0019828 | ☐ | ☐ | 1.2281 |

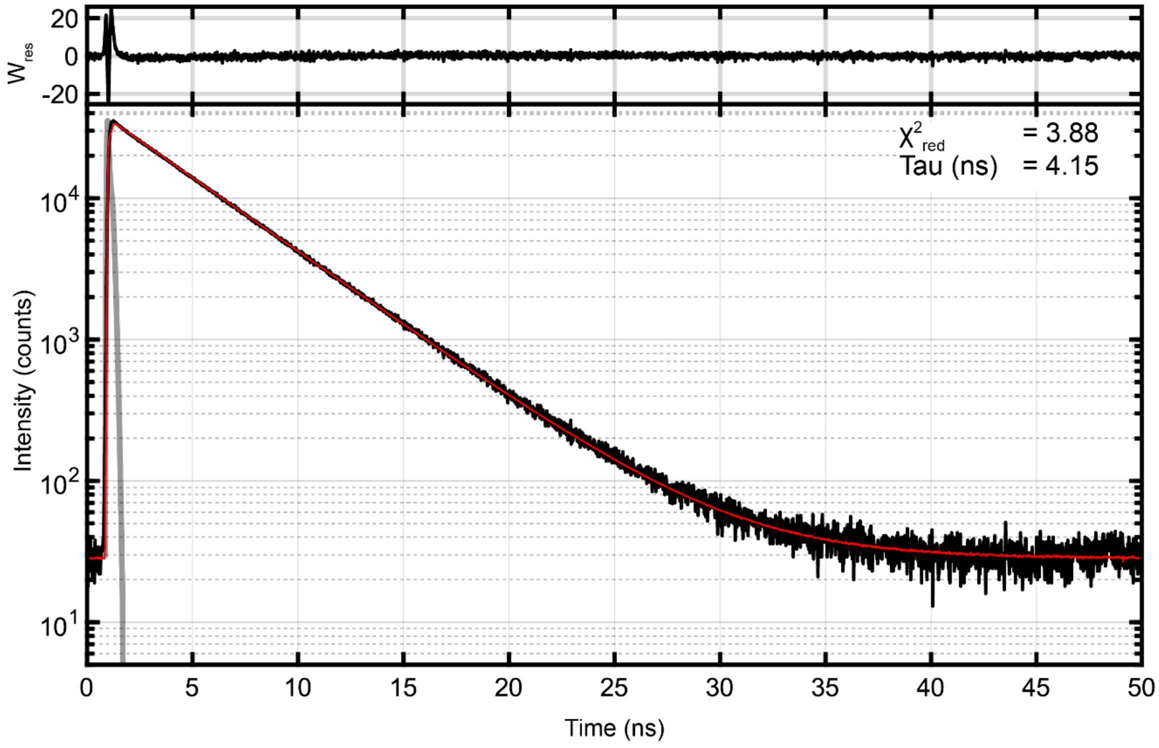


Figure S5: Illustration of PAM-*FCSfit* and PAM-*Taufit* data from PAM to verify microscope alignment and characterize reference lifetime for phasor referencing. (TOP) Fluorescence correlation spectroscopy (FCS) analysis with ATTO-488and ATTO-647N on our homebuilt confocal results in excellent focal parameters and photon sensitivity and is performed every measuring session. (BOTTOM) Reconvolution data fitting with loaded IRF to identify the lifetime of the measured reference sample used in phasor calculation.

#### Supplemental Figure 6

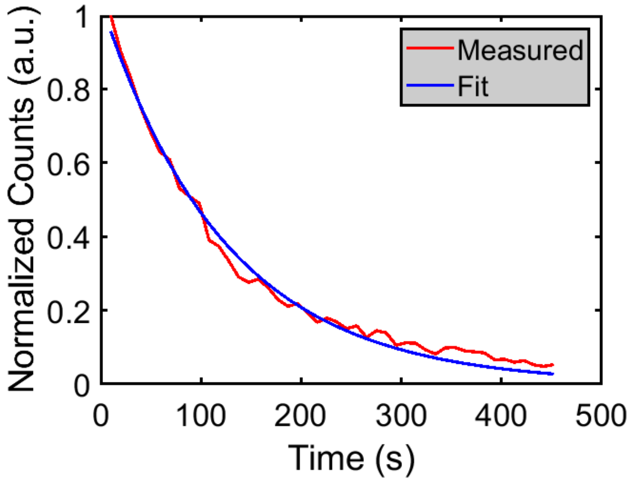


Figure S6 Exemplary plot demonstrating the exponential fit on a single HIV-1 particle evaluated for its fluorescence output over time containing IN-eGFP. Bleaching rates are derived from the exponential fit.

### Supplemental Figure 7

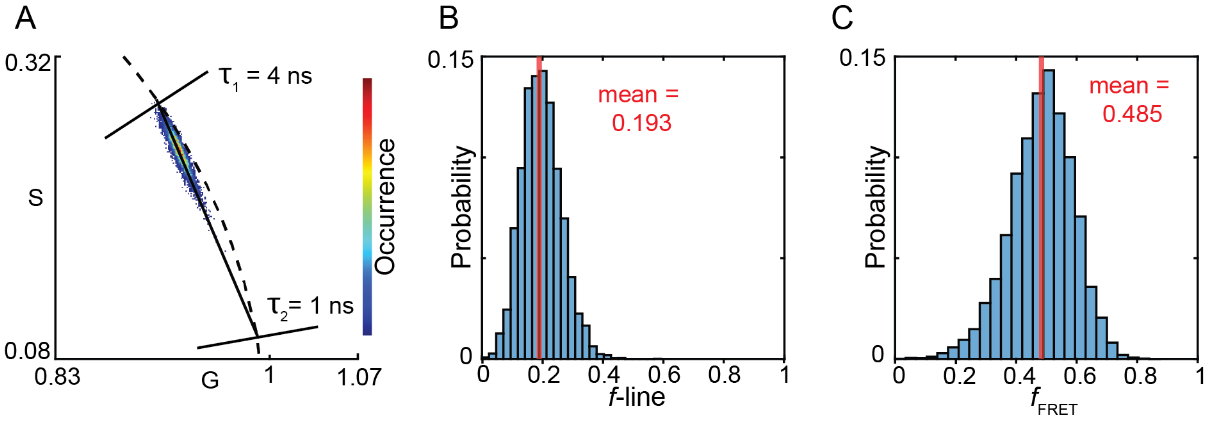


Figure S7: Simulation data illustrating the determination of concentration fraction. (A) Phasor plot resulting of a mixture of two fluorescent species with lifetimes of 4 ns and 1 ns for which the latter has an intensity of 1/4$^{th}$ of the 4 ns component. Species are simulated to have free yet identical diffusion so to obtain an image of simulated photon data containing pixels with a distribution near 50/50 concentration. Similarly, FRET experiments often contain pixels with unquenched and quenched forms of the donor. A fraction line (black line) can be drawn in between. (B) All phasor positions can be summed to the closest position on the fraction line leading to a distribution of fractions. (C) Using Eq. 20 (see also supplemental note SN1), the intensity fractions can be converted to concentration fractions resolving the simulated 1:1 ratio of species.

#### Supplemental Figure 8

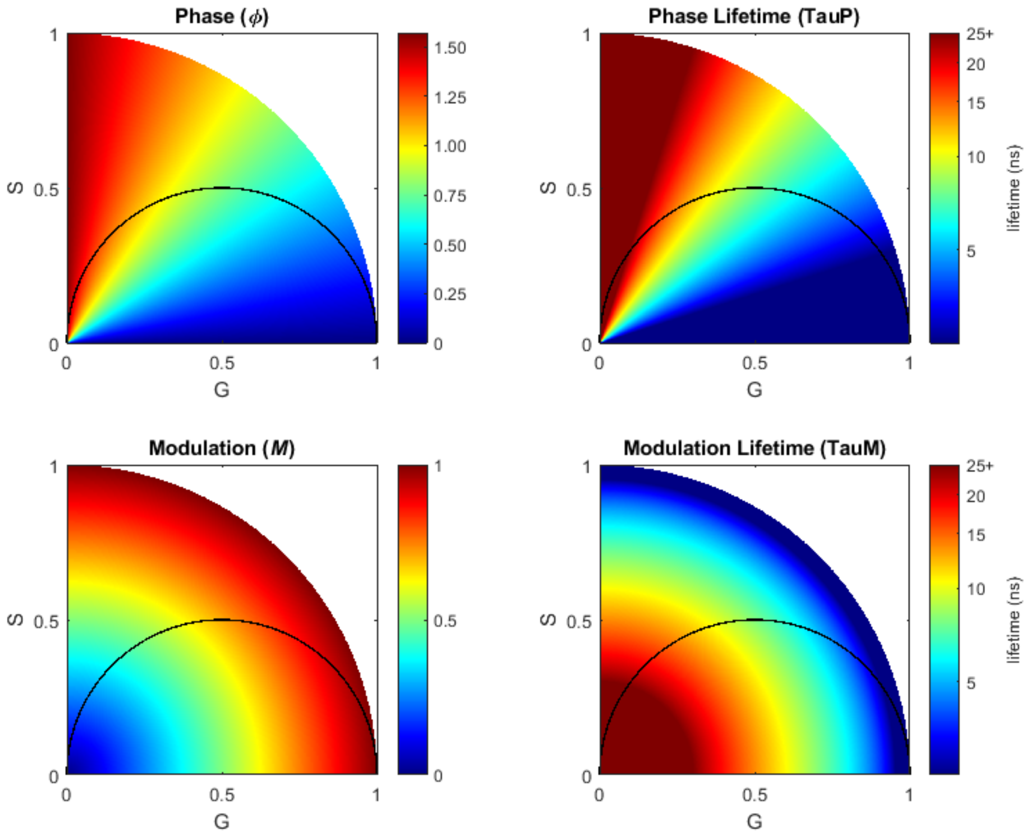


Figure S8: Color-coding of phase and modulation values in the first quadrant of the universal circle, containing the semi-circle. While the Phase and Modulation are clearly different, the therefrom calculated phase lifetime and modulation lifetime display equal values along the crosscut of the colored quadrant and the semicircle. This demonstrates the phasor property that mono-exponential lifetimes situated on the semicircle have an identical phase and modulation lifetime (and only there).

### Supplemental Figure 9

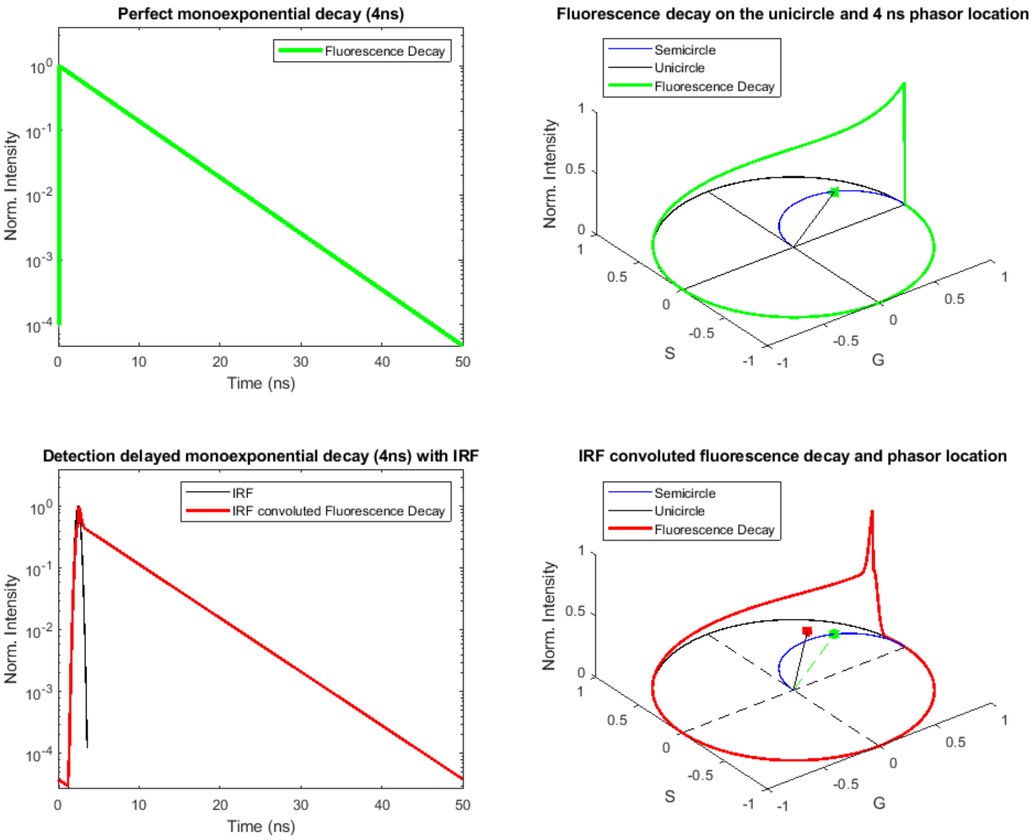


Figure S9: Graphical representation of the effect of IRF convolution in a perfect decay (4 ns). Green represents a prefect decay and a visual representation on the universal circle (unicircle) of that decay, leading to a phasor spot perfectly on the expected location on the semi-circle. (B) When an IRF peak is convoluted with the decay, and there is a delay of detection on the TCSPC, the resulting phasor location is no longer on the semi-circle due to the added non-exponential component. This effect is corrected by applying a phase and modulation change extracted from a reference sample.

### Supplemental Figure 10

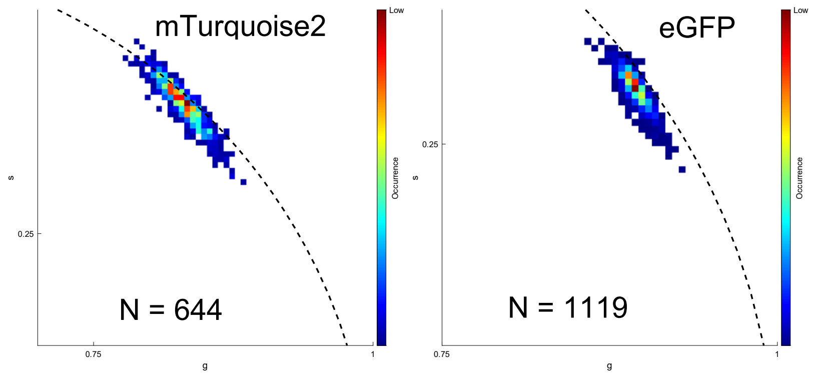


Figure S10: Donor-only phasors of HIV-1 IN FRET particles used to determine FRET trajectory starting point (DO) in Fig. 4.

#### Supplemental Figure 11

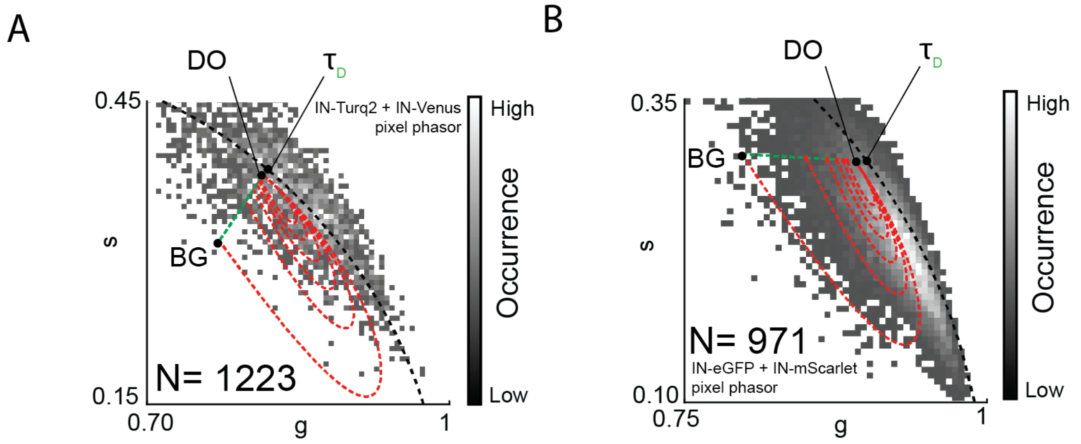


Figure S11: Pixel based phasor of the measured HIV-1 IN FRET particles with IN-mTurquoise2 and IN-mVenus (A) or IN-eGFP and IN-Scarlet (B). As can be seen the trajectories for 0-50% passive donor fractions (dashed red lines, larges trajectory is 0% passive donors, smallest is 50%) are difficult to match with measured pixel data. A pixel selection threshold of 20 photons/pixel was used to exclude non-particle pixels.
